# Organic Germanium (Ge-132) reduces glycative damage while maintaining cellular stress signaling: evidence of functional dissociation

**DOI:** 10.64898/2026.04.22.720084

**Authors:** Alejandro Ponce Mora, Yasin Fauzi El-Adhiri, Gabrielle Guillamin, Amanda Martell Vergara, Antonella Locascio

## Abstract

Organic germanium, particularly carboxyethyl germanium sesquioxide (Ge-132), has been investigated for decades in relation to diverse biological effects, with a strong emphasis on its antioxidant properties. However, the available literature remains dispersed, encompassing heterogeneous experimental models and endpoints that limit mechanistic interpretation. While antiglycative activity has been described at the biochemical level, its downstream gene regulatory consequences under glycative stress remain inconsistently characterized.

Here, we combined systematic review of the literature of experimental studies with targeted molecular analysis in a standardized cellular model. The literature mapping was used to guide pathway selection rather than to establish quantitative associations. Based on patterns emerging from literature, we focused on pathways associated with glycative stress responses, including carbonyl stress, inflammatory signaling, and autophagy regulation. Gene expression analysis revealed a limited and selective modulation of regulatory pathways under glycative stress conditions, consistent with a context-dependent effect rather than broad transcriptional reprogramming. In parallel, protein analysis showed reduced intracellular accumulation of advanced glycation end products (AGEs) in Ge-132–treated cells under glycative stress conditions.

Importantly, these findings support a dissociation between glycative damage reduction and cellular stress-response pathways. This combined approach helps interpretation of previously fragmented observations across the literature and highlights gene regulation under glycative stress as a relevant but still unresolved aspect of organogermanium biology.

## 1. Introduction

Germanium (Ge) is a naturally occurring trace element found in soils, rocks, and plants such as garlic, ginseng, aloe, shiitake mushrooms, and pearl barley [1,2]. It has a face-centered cubic lattice, and physicochemical properties intermediate between metals and non-metals, supporting diverse technological and biomedical uses [1,2]. Initially used in electronics, interest in its biological properties grew in the 1970s–1980s. Germanium compounds are classified as inorganic or organic, differing in physicochemical and biological properties. Inorganic forms are poorly soluble, accumulate in organs, and can cause dose-dependent, often irreversible toxicity, including nephrotoxicity and neurotoxicity [2–7]. Nevertheless, germanium-68 (⁶⁸Ge) is used in nuclear medicine as a parent radionuclide for gallium-68 (⁶⁸Ga) in PET imaging for cancer and inflammation detection [8–10].

Organic germanium compounds show distinct biological and toxicological profiles. The most studied is Ge-132 (Repagermanium), a water-soluble polymer hydrolyzing to 3-(Trihydroxygermyl) propanoic acid (THGP), that exhibit low toxicity and minimal tissue accumulation [3,11–16]. THGP forms complexes with cis-diol biomolecules, including saccharides and nucleotides, underlying its biological effects, particularly modulation of oxidative stress [17–19].

Ge-132 has been associated with effects on mitochondrial metabolism, immune signaling, and stress adaptation. It modulates oxidative stress not by direct ROS scavenging but by regulating mitochondrial metabolism and antioxidant systems, restoring redox homeostasis [7,20,21]. THGP preserves ATP levels, promotes oxidative phosphorylation, and limits ROS and lipid peroxidation through multiple mechanisms, including complex formation with biomolecules, inhibition of adenosine deaminase, stimulation of aldehyde reductase, and inhibition of AGE formation [2,7,21]. It has also been reported to modulate transcriptional responses, suppressing *NR4A2* and inflammatory mediators, such as *IL-6* and *CXCL2*, and regulating apoptosis-related pathways including KEAP1–BAX–caspase-3 and *BCL-2* [22,23]. However, the high variability in models and conditions used in all those reports complicates interpretation and generalization of these studies.

Ge-132 also influences immune and inflammatory responses, enhancing IFN-γ production, NK cell activity, and B-cell responses [19,24–27]. For instance, THGP regulates inflammation by sequestering extracellular ATP and inhibiting caspase-1–mediated IL-1β release [24,28]. It also promotes tissue repair via immune cell recruitment and *TGF-β*–mediated collagen synthesis [29], and preservation of bone integrity [30].

Antitumor effects have been also reported, including reduced tumor viability, inhibition of metastasis, modulation of SIRP-α–CD47 signaling, and induction of cell-cycle arrest and apoptosis [25,26,31–36]. Clinical observations suggest additional supportive effects in palliative care [37,38].

Similar protective roles have been observed in plants, where germanium enhances growth and salt stress tolerance via oxidative balance modulation [39–43].

*In-vivo* and cellular models, Ge-132 also affect aging-related pathways by supporting antioxidant defenses, immune function, and cell renewal. It has been documented that THGP enhances clearance of senescent Red Blood Cells and reduces oxidative stress–induced apoptosis, with associated modulation of *Nrf-2* and *Bax* genes, while preserving mitochondrial function and ATP levels [31,44–46].

The antioxidant properties of Ge-132 and THGP extend to the modulation of glycative stress. Rather than reflecting a classical antioxidant mechanism, this effect appears to be mediated by complex formation with cis-diol–containing monosaccharides, thereby interfering with the Maillard reaction and reducing the formation of AGEs [47–51]. This antiglycative activity has been demonstrated in diabetic experimental models, where protein glycation is significantly inhibited [52–55]. In contrast to the relatively well-described biochemical antiglycative effects, the downstream gene regulatory responses associated with glycative stress under Ge-132 exposure remain largely unexplored. This gap limits current mechanistic understanding, particularly considering that AGEs formation is a major source of reactive oxygen species and that its inhibition may indirectly contribute to the regulation of oxidative stress and maintenance of cellular homeostasis.

Extending germanium described properties, the intestinal microbiota represents an additional level at which dietary Ge-132 may influence host homeostasis. Although Ge-132 does not appear to markedly alter caecal microbial composition, it interacts functionally with the microbial ecosystem, particularly in the presence of prebiotics or probiotics [17,28,56]. In combination with oligosaccharides such as raffinose and with *Lactobacillus* species, Ge-132 enhances mucosal immune responses [28,57]. Interestingly, Ge-132 alone has been reported to increase β-glucuronidase activity, a marker associated with elevated colon cancer risk, although this effect is abolished when prebiotics are co-administered, underscoring the context-dependent biological effects of Ge-132 [56].

In short, these observations suggest that organic germanium influences multiple biological processes, including redox regulation, immune signaling, metabolic adaptation, and stress resilience. However, the available evidence originates from heterogeneous experimental systems and addresses distinct biological endpoints, resulting in a fragmented body of literature that limits mechanistic interpretation and precludes meaningful quantitative synthesis.

Rather than aiming to perform a quantitative synthesis, the present study adopts an evidence-informed experimental approach, in which systematic review of the literature is used not to broaden the scope of the work, but to provide a framework for interpreting mechanistic inconsistencies across literature. This is particularly relevant in the context of glycative stress, where reductions in AGE accumulation have been attributed to diverse and not always convergent mechanisms, including direct inhibition of glycation reactions through interaction with cis-diol–containing molecules [47,51], as well as indirect effects mediated by modulation of oxidative stress and cellular signaling pathways [20,22,58].These interpretations are often proposed independently, without a clear mechanistic integration between biochemical and cellular levels. Therefore, glycative stress is used here as a focused experimental entry point to address a broader conceptual question: whether the biological effects attributed to Ge-132 reflect coordinated cellular regulatory mechanisms or are primarily driven by upstream biochemical interactions (Figure 1).

**Figure 1.**
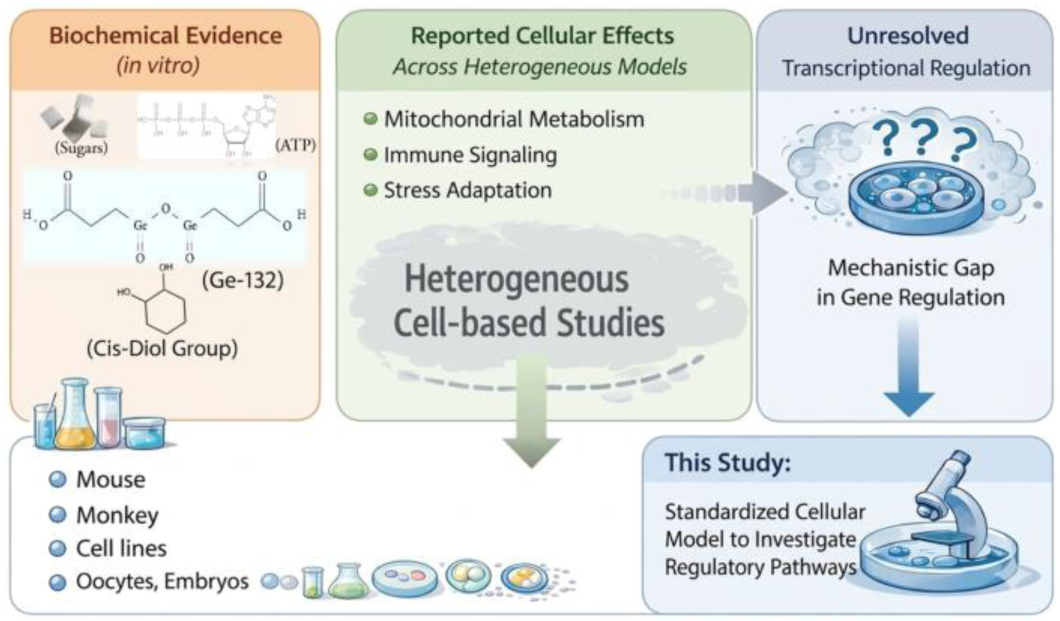
Conceptual overview of current knowledge on organic germanium (Ge-132). While biochemical studies show its interaction with cis-diol–containing molecules, cellular studies report diverse effects including mitochondrial metabolism, immune signaling, and stress responses. The mechanisms linking these observations remain unclear, particularly at the transcriptional level. Here, we address this gap by examining the antiglycative activity of Ge-132 in a standardized cellular model.

## 2. Results

### 2.1. Systematic mapping of the literature

#### 2.1.1. Screening and selection

The literature search and screening process is summarized in Figure 2. After removal of duplicates and application of predefined inclusion criteria, 24 studies investigating the biological effects of organic germanium (Ge-132) were selected for qualitative evaluation. This dataset represents a subset of currently available experimental evidence based on the applied criteria addressing the biological activity of Ge-132.

**Figure 2.**
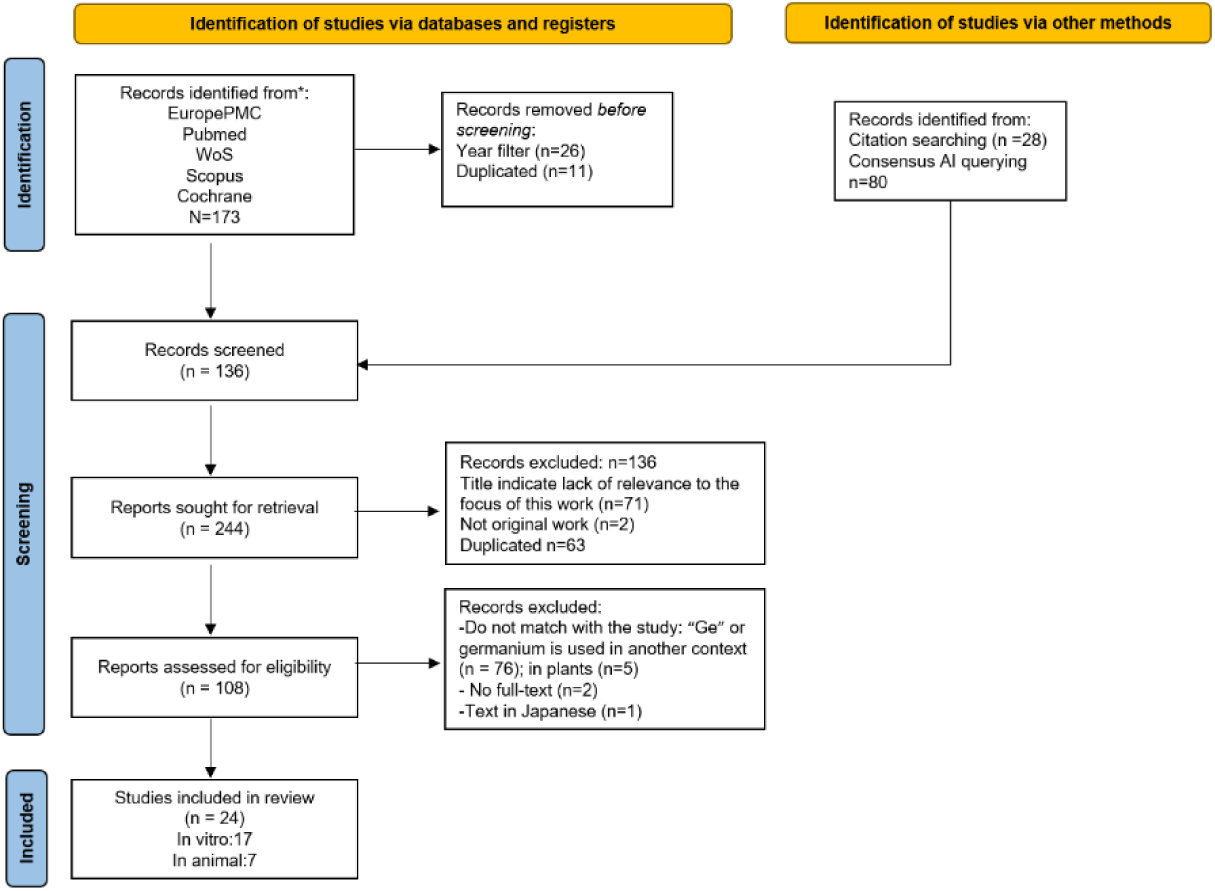
PRISMA flow diagram showing the process of selection of evidences published in the last 15-years.

The studies were evaluated by comparing those conducted exclusively *in-vitro* with those performed in *in-vivo* models. Only a small number of studies combined both *in-vitro* and *in-vivo* approaches. The analysis of evidences identified three main biological axes: redox regulation, immune/inflammatory signaling, and glycative/carbonyl stress (Figure 3a). Among those, redox and immune-related pathways were the most frequently investigated, whereas glycative stress was comparatively underrepresented. This distribution indicates that, although multiple biological effects have been attributed to Ge-132, research efforts have predominantly focused on classical stress and immune pathways, while glycative stress, despite its relevance in metabolic and age-related processes remains insufficiently explored.

**Figure 3.**
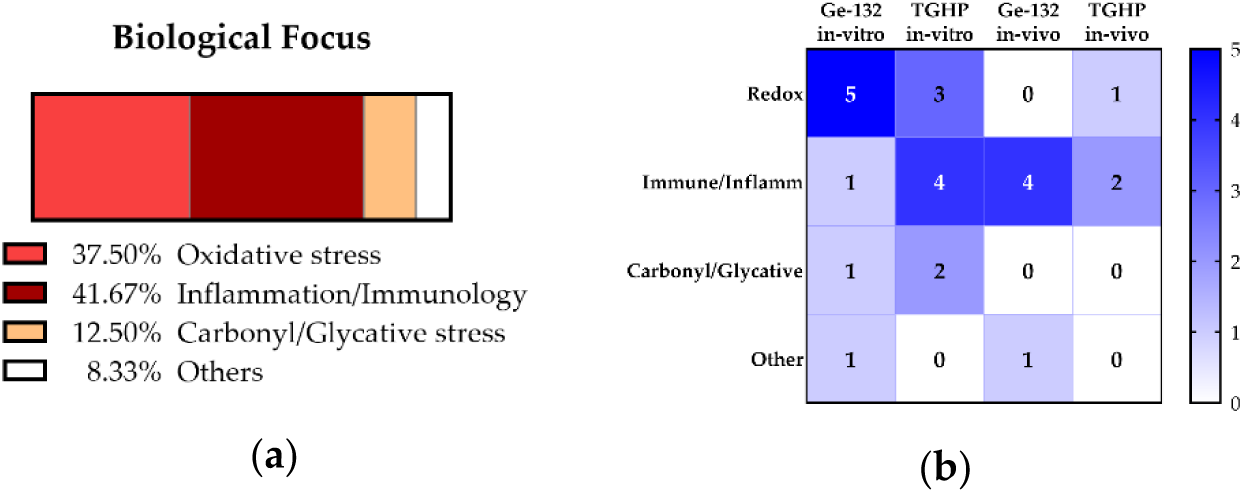
Evidence mapping. (a) Distribution of experimental studies on organogermanium biological activity by area of investigation, over the past 15 years. (b) Distribution of experimental evidence across experimental models (*in-vivo, in-vitro*), compound tested (Ge-132 or THGP) and outcome domain (redox, immune/inflammatory, glycative stress, and other). Color intensity indicates the number of studies per category.

Evidence mapping integrating compound type, experimental model, and outcome domain confirmed this imbalance (Figure 3b). A predominance of *in-vitro* studies and redox-related outcomes was observed across both Ge-132 and THGP. In contrast, glycative stress studies were limited and mainly restricted to *in-vitro* systems.

Overall, these observations highlight a scattered evidence base, with limited mechanistic continuity and a lack of study addressing gene regulatory responses under glycative stress conditions.

#### 2.1.2 Implications for quantitative revision

The studies included in this analysis encompass a wide range of biological processes, including redox regulation, immune signaling, apoptosis, and metabolic modulation. However, substantial heterogeneity in experimental models, exposure conditions, and study design limits direct comparability across studies and precludes meaningful quantitative mapping (Tables 1 and 2). This heterogeneity is reflected not only in the diversity of biological endpoints but also in the variability of experimental systems, compound forms, and dosing strategies. As a result, the available evidence cannot be integrated into a unified quantitative framework, reinforcing the need for standardized experimental approaches to enable mechanistic interpretation.

**Table 1.** Overview of *in-vitro* studies using Ge-132 in clinical research.

| Authors | Study area | Population model | Dose exposure |
| --- | --- | --- | --- |
| Nakamura *et al.*, 2015[59] | Oxidative stress, Immunity, Inflammation | Human Keratinocytes | THGPA 0.25 to 500 mmol/l. THGPA 0, 2.5, and 5 mM for adrenaline; 0, 1, and 10 mM for ATP. |
| Kim *et al.* 2015[23] | Reproductive biology, Oxidative stress, Apoptosis | Porcine Oocytes | 100 mg/mL, 200 mg/mL, and 400 mg/mL Ge-132. |
| Shimada *et al.*, 2015[60] | Biochemistry, Immunology, Diabetes Research | *In-vitro* (water solution) | 0.5 M for THGPNa |
| Tezuka *et al.*, 2017[20] | Oxidative stress, Oncology, Inflammation | Hepatocytes from Japanese macaque | 0, 0.001, 0.01, 0.1, or 1 μM Ge-132 |
| Kim *et al.*, 2017[61] | Oxidative stress, Apoptosis, Cell signaling (KEAP1) | Porcine IVF embryos | 0, 100, 200, and 400 μg/ml Ge-132 |
| Shimada *et al.*, 2018 [62] | Biochemistry, Inflammation, Pain management | *In-vitro* (water solution) | THGP: 0 -100 mM |
| Wada *et al.*, 2018 [46] | Oxidative stress, Cell proliferation, Immunology, Oncology | Mammalian ovary cells; Chinese Hamster Ovary (CHO-K1) cells; HeLa cells, SH-SY5Y cells | 0- 5 mM Ge-132 |
| Filonova *et al.*, 2019[63] | Antioxidant properties | Mouse different tumor cells; ovary cells | NMR spectroscopy: Ge-132 at $1.3 \times 10^{-1}$ M and $1.3 \times 10^{-2}$ M. Cyclic voltammetry: Ge-132 at $3 \times 10^{-3}$ M, $6 \times 10^{-3}$ M, and $9 \times 10^{-3}$ M |
| Takeda *et al.*, 2019 [22] | Oxidative stress, Cell signaling and inflammatory response | Human dermal fibroblasts (NHDFs) | 0, 5.9, 59, 590, and 5900 μM THGP.LC MS/MS analysis: 0, 1, and 10 mM THGP. PI/Hoechst staining: 0, 5.9 mM THGP. |
| Wang *et al.*, 2020 [58] | InflammationImmunity, Cell Signaling (NF-κB and MAPK pathways) | Primary mouse Mammary Epithelial Cells | Ge-132 at 0.5 μg/mL, 1 μg/mL, and 2 μg/mL |
| Mertens *et al.*, 2020 [64] | Oncology, Oxidative Stress | MDA-MD-175; MDA-MB-231; MRC5; H460; A2780 | Germanium(IV)-diketonat at 5 μM or 10 μM |
| Baidya *et al.*, 2021 [65] | Immunity, Antiviral strategies, Inflammation | RAW 264.7; HEK293T; A549; MDCK | THGP at 0, 20, 200, and 2000 μg/mL |
| Azumi *et al.*, 2022 [28] | Inflammation | Human THP-1 monocytes; mouse RAW264 | THGP at 0.05, 0.5, 1.2, 5 and 12 mM; Ge-132 500 mM |
| Sekiguchi *et al.*, 2023 [66] | Pain management, Oxidative stress, Calcium Channel Signaling | Cav3.2-transfected HEK_293 (human kidney) | THGP at 1, 3 and 10 mM |
| Azumi *et al.*, 2023 [24] | Immunity, Oncology | RAW 264.7 mouse macrophage-like cells | 500 μM THGP. Dose dependent: THGP from 50 to 5000 μM |
| Cruz *et al.*, 2024[67] | Oxidative stress | Bovine Sperm cells | 0-500-1000 μg/mL of Ge-132 |
| Takeda *et al.*, 2024 [44] | Immunology, Antioxidant, Cardiovascular/<br>Gastrointestinal health | Mouse cell RAW264.7 | 0, 50, 500, and 5000 µM THGP |

Abbreviations: Ge-132, carboxyethyl germanium sesquioxide; THGP, 3-(trihydroxygermyl)propanoic acid. Study area refers to the primary biological process investigated in each study, as defined by the authors. Population models include *in-vitro* cellular systems. Dose/exposure is reported as described in the original publications.

**Table 2.**
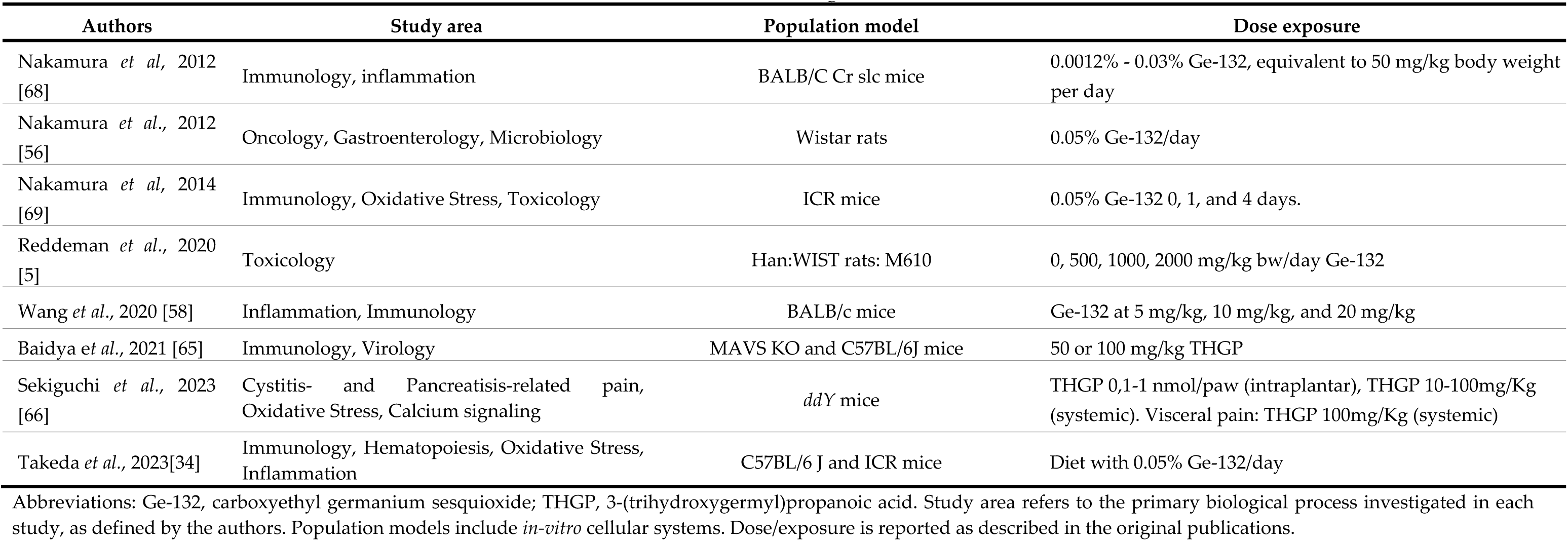
Overview of *in-vivo* studies using Ge-132 in clinical research.

| Authors | Study area | Population model | Dose exposure |
| --- | --- | --- | --- |
| Nakamura *et al.*, 2012 [68] | Immunology, inflammation | BALB/C Cr slc mice | 0.0012% - 0.03% Ge-132, equivalent to 50 mg/kg body weight per day |
| Nakamura *et al.*, 2012 [56] | Oncology, Gastroenterology, Microbiology | Wistar rats | 0.05% Ge-132/day |
| Nakamura *et al.*, 2014 [69] | Immunology, Oxidative Stress, Toxicology | ICR mice | 0.05% Ge-132 0, 1, and 4 days. |
| Reddeman *et al.*, 2020 [5] | Toxicology | Han:WIST rats: M610 | 0, 500, 1000, 2000 mg/kg bw/day Ge-132 |
| Wang *et al.*, 2020 [58] | Inflammation, Immunology | BALB/c mice | Ge-132 at 5 mg/kg, 10 mg/kg, and 20 mg/kg |
| Baidya *et al.*, 2021 [65] | Immunology, Virology | MAVS KO and C57BL/6J mice | 50 or 100 mg/kg THGP |
| Sekiguchi *et al.*, 2023 [66] | Cystitis- and Pancreatitis-related pain, Oxidative Stress, Calcium signaling | *ddY* mice | THGP 0,1-1 nmol/paw (intraplantar), THGP 10-100mg/Kg (systemic). Visceral pain: THGP 100mg/Kg (systemic) |
| Takeda *et al.*, 2023[34] | Immunology, Hematopoiesis, Oxidative Stress, Inflammation | C57BL/6 J and ICR mice | Diet with 0.05% Ge-132/day |

Abbreviations: Ge-132, carboxyethyl germanium sesquioxide; THGP, 3-(trihydroxygermyl)propanoic acid. Study area refers to the primary biological process investigated in each study, as defined by the authors. Population models include *in-vitro* cellular systems. Dose/exposure is reported as described in the original publications.

Observation of this fragmented literature highlights a gap particularly at the level of mechanistic and gene regulatory responses associated with glycative stress. Guided by these findings, we next investigated whether the antiglycative effects described at the biochemical level are associated with measurable gene regulatory responses under controlled glycative stress conditions.

### 2.2 Risk of Bias Assessment

To evaluate methodological reliability, each study was assessed using a CASP-adapted risk-of-bias framework [70].

In the case of *in-vitro* studies (Figure 4a), the RoB assessment revealed moderate methodological variability. Sequence generation procedures were frequently insufficiently reported, resulting in a substantial proportion of unclear risk classifications, whereas allocation concealment was generally well described. Incomplete outcome data showed a balanced distribution between low and unclear risk, suggesting acceptable reporting despite limited documentation of exclusions. Notably, selective outcome reporting was predominantly classified as unclear, indicating that experimental endpoints were often not predefined or fully described. Additionally, the presence of high-risk classifications within the “Other sources of bias” domain primarily reflects the involvement of research groups associated with the development or provision of the organogermanium compound under investigation. While such collaborations may facilitate compound standardization and experimental consistency, the limited independent replication across laboratories may introduce potential sponsorship-related bias.

**Figure 4.**
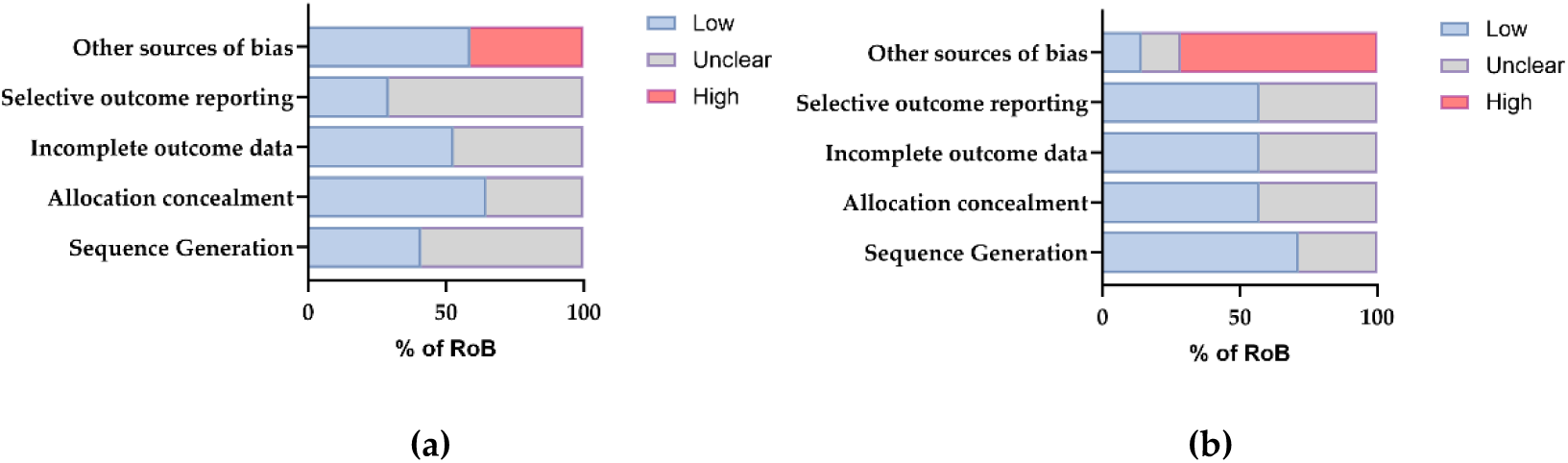
Risk-of-bias assessment of methodological reliability across quality domains. Data represent the percentage of risk in the total of publications included in each sample category. (a) *In-vitro* based research; (b) *In-vivo* based research.

The RoB assessment for *in-vivo* studies (Figure 4b) indicated a generally adequate experimental structure, with low risk predominating in the domains of sequence generation and incomplete outcome data. Allocation concealment and selective outcome reporting were frequently classified as unclear due to incomplete methodological reporting. The “Other sources of bias” domain showed a predominance of high-risk classifications, reflecting limited independent replication across research groups and potential structural influences related to compound sourcing.

Overall, the comparison between *in-vitro* and *in-vivo* studies highlights substantial variability in methodological reporting and experimental design across the Ge-132 literature, supporting the need for standardized experimental frameworks to enable coherent mechanistic interpretation. An additional feature emerging from the literature analysis is the apparent lack of continuity across research lines. Most studies represent isolated investigations that are not followed by subsequent work exploring the same biological mechanisms in greater depth or across complementary models. While some degree of thematic progression can be observed within specific research groups, particularly in studies originating from Asian research institutes. However, these efforts tend to shift across different biological contexts rather than systematically advancing a single mechanistic pathway. This pattern contributes to the fragmentation of the field and further limits the consolidation of robust and reproducible mechanistic frameworks.

### 2.3 Cytotoxicity assay across distinct cell lines

Three different cellular models were initially evaluated to assess Ge-132 cytotoxicity and differential sensitivity: human lens epithelial cells (HLECs), representing a system susceptible to glycation-related protein damage; ARPE-19 cells, as a model of metabolically active retinal tissue exposed to oxidative stress; and mouse embryonic fibroblasts (MEFs), a well-established system for studying redox regulation and stress-response signaling [71–75]. The concentration range of Ge-132 was selected based on the doses most frequently reported in the literature (Table 1).

In general, we observe that Ge-132 did not induce relevant cytotoxicity within the concentration range used for ARPE-19 and HLEC cells. However, MEFs displayed a higher sensitivity to Ge-132 at elevated concentration (Figure 5). MEFs were therefore selected for subsequent mechanistic analyses as they provide a suitable dynamic range to detect stress-related and modulatory effects. The use of a single, consistent cellular context in the present study enables direct comparison across multiple biological pathways within the same experimental framework, reducing variability associated with cross-model comparisons and facilitating mechanistic interpretation.

**Figure 5.**
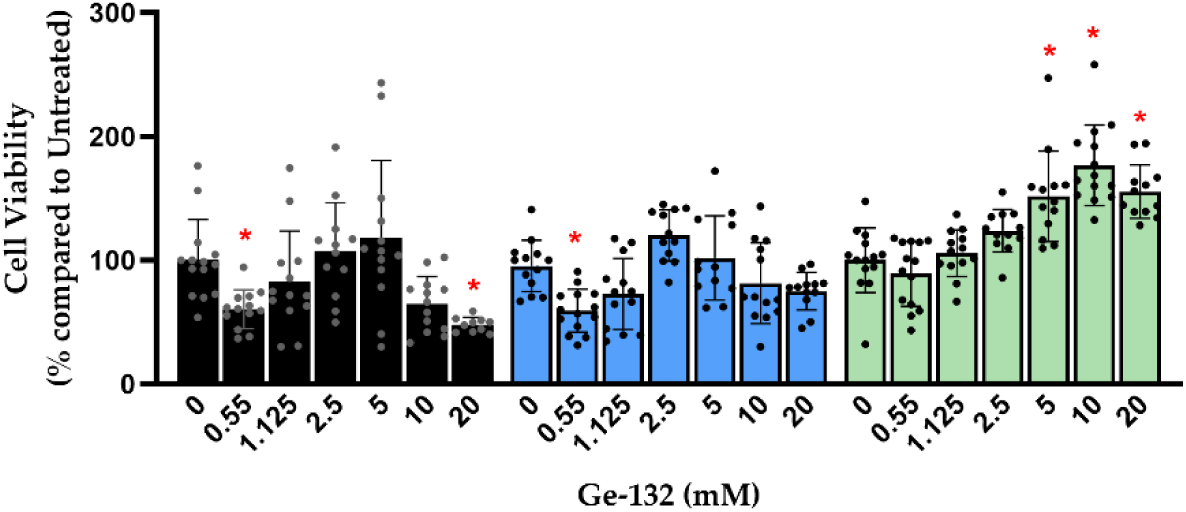
Cytotoxicity assessment of Ge-132 across different cell lines. Cell viability assessed in MEFs (black bars), HLECs (blue-bars), and ARPE-19 cells (green-bars) following exposure to increasing concentrations of Ge-132. Data represent mean of five biological replicates, with 10 to 15 technical replicates ± SD. Significant differences were marked with a red *, after applying a one-way ANOVA test followed by Dunnett’s multiple comparison correction of each treatment *vs* the corresponding control.

### 2.4 Antiglycative activity of Ge-132

Based on the gaps identified in the literature mapping, we focused our experimental analysis on transcriptional responses associated with glycative stress.

To evaluate the antiglycative activity of Ge-132 across different levels of biological complexity, we first assessed its effect in a cell-free Maillard reaction system using concentrations equivalent to those applied in cytotoxicity assays across the three cellular models. Under these conditions, Ge-132 reduced the formation of glycation products, detectable from the concentration of 5 mM onwards, with a dose-dependent effect (Figure 6). These data are consistent with the antiglycative activity of Ge-132, as previously described in literature, highlighting the reduction of AGE accumulation even under glycative stress conditions.

**Figure 6.**
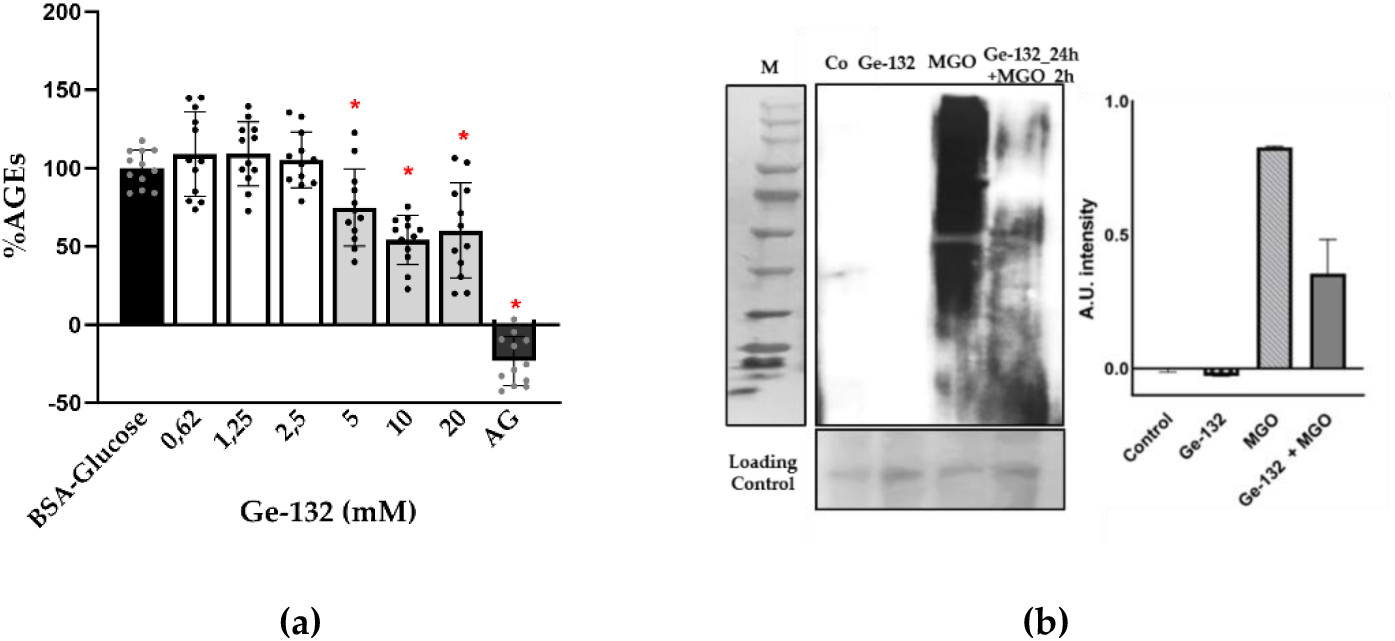
Antiglycative activity of Ge-132 in cell-free and cellular systems. (a) *In-vitro* Maillard reaction assay showing dose-dependent reduction of AGE formation in the presence of Ge-132 (5–20 mM). AGE levels are expressed relative to the positive control (BSA+Glucose). Aminoguanidine (AG) was used as a reference inhibitor of AGE formation. Graph is representative of three experimental replicates, including 12 technical replicates. Multi-group comparisons were analyzed using one-way ANOVA followed by Dunnett’s post hoc test. Error bar represents SD. (b) Immuno-detection of methylglyoxal-derived protein adducts in MEF cells exposed to MGO, with or without Ge-132 pre-treatment. α-MG was used to detect MGO-derived protein adducts. Ponceau staining was used as a loading and membrane enrichment control. Densitometric quantification of the immune-localization signal is shown.

We next investigated whether this effect could be reproduced in a cellular context. Mouse embryonic fibroblasts (MEFs) were exposed to glycative stress induced by methylglyoxal (MGO), in the presence or absence of Ge-132. In addition to co-treatment conditions, and consistent with previously reported short-term glycative stress models [76], a sequential exposure of MGO for 2 hours followed the treatment with Ge-132. This setting was included to explore potential preventive or modulatory effects of GE-132 in case of glycative insult. Western blot analysis revealed an accumulation of AGEs-modified proteins under MGO treatment, consistent with the establishment of glycative stress conditions. Ge-132 significantly reduced intracellular AGEs accumulation in fibroblasts exposed to methylglyoxal, particularly when administered as pre-treatment. Importantly, Ge-132 alone did not modify AGEs levels respect the untreated control, indicating that its effect is conditional upon stress exposure rather than constitutive inhibition of glycation reactions (Figure 6b). These findings indicate that the anti-glycative activity previously described in solution-based systems can be reproduced in a cellular context, supporting the ability of Ge-132 to modulate glycative stress under biologically relevant conditions.

### 2.5 Gene expression analysis

To investigate whether the anti-glycative effects described at the biochemical level, and confirmed here by Western blot analysis, are associated with measurable cellular responses, we analyzed the expression of selected regulatory genes participating in pathways related to glycative stress, in our standardized *in-vitro* MEFs model. The selection of target genes was based on their known established roles as key regulatory nodes within pathways associated with glycative stress in the literature [77–82].

#### 2.5.1 Antioxidant response

Multiple *in-vitro* studies have described antioxidant-like effects associated with Ge-132 without demonstrating strong direct ROS scavenging, suggesting instead indirect modulation of redox signaling pathways (see Table 1 and 2). Animal studies similarly reported reduced oxidative cell damages and improved physiological outcomes following Ge-132 administration.

To evaluate the impact of glycative stress on oxidative stress–related transcriptional responses, we analyzed the expression of a key gene involved in the activation of this pathway, specifically *NRF2*. Related to *NRF2*, we studied the expression of downstream direct targets as HMOX1 and NQO1, selected as central regulators of redox adaptation (Figure 7a). Additional NRF2-related targets involved in glutathione synthesis (*GCLC* and *GCLM*) are included.

**Figure 7.**
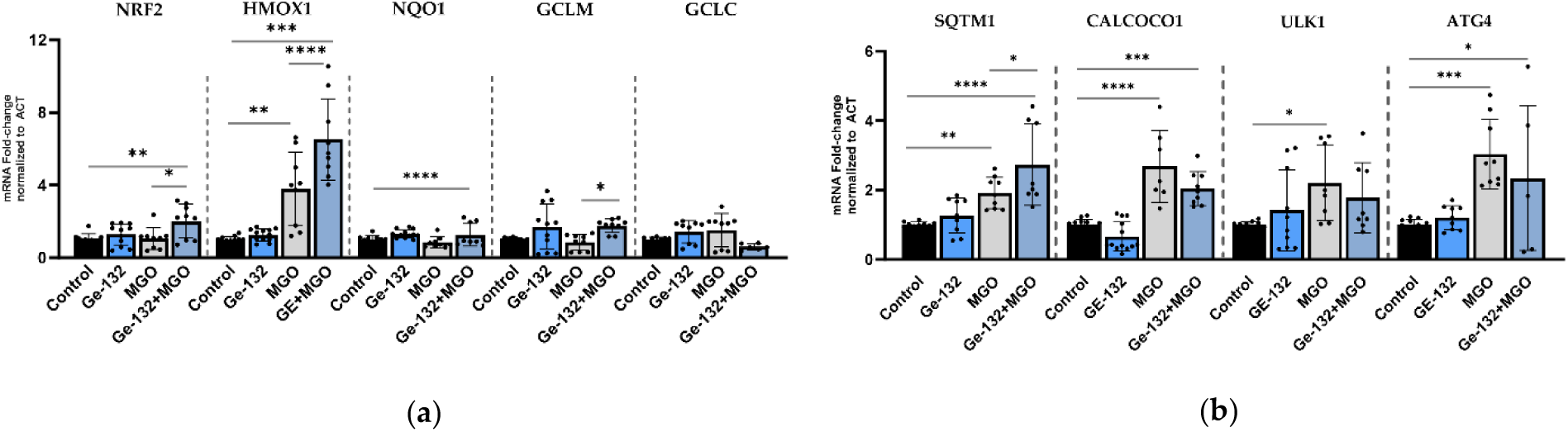
Transcriptional response of antioxidant and autophagy-related genes under glycative stress and Ge-132 treatment. (a) Relative expression of *NRF2* pathway genes (*NRF2, HMOX1, NQO1*). MGO induced upregulation of HMOX1. Ge-132 alone produced no significant changes, while combined Ge-132+MGO treatment induced modest expression increase of *NRF2* and NRF2-targets. *GCLC* and *GCLM* expression levels are shown as additional NRF2-related targets and did not show relevant changes across conditions. (b) Relative expression of autophagy-related genes. MGO induced upregulation of cargo-recognition receptors (*SQSTM1, CALCOCO1*) and early regulatory components (*ULK1, ATG4*). Data are normalized to actin and presented as mean of three biological replicates ± SD. Student’s t-test was used for statistical significance comparing each individual treatment *vs* related control, significance is indicated.

MGO alone did not significantly induce *NRF2* transcription, whereas combined Ge-132+MGO treatment resulted in a modest but consistent increase. *NQO1* is not sensitive to MGO treatment neither when germanium is applied. *HMOX1*, which is a recognized downstream target of NRF2, was significantly upregulated. This result indicates activation of a cellular oxidative stress response. Although *HMOX1* is a canonical *NRF2* target is not exclusively dependent on *NRF2,* being also regulated by multiple stress-responsive transcription factors, including *NF-κB*, which can drive its induction under inflammatory or oxidative conditions [83].

Ge-132 treatment did not induce significant changes in these key antioxidant genes. On the other hand, under combined Ge-132 and MGO treatment, both *NRF2, HMOX1* and *NQO1* expression were induced, consistent with activation of the oxidative stress response is somehow preserved. *GCLC* and *GCLM* showed no significant changes across experimental conditions, consistent with a selective activation of antioxidant responses.

Although transcript levels do not necessarily match protein abundance, and basal protein activity may depend on protein stability or turnover, the MGO-induced increase in *NRF2* transcription in the presence of Ge-132, still indicates activation of oxidative stress signaling. Since Ge-132 alone did not elicit a comparable transcriptional response, these findings together suggest that Ge-132 may mitigate glycative damage primarily through a mechanism likely mediated by upstream biochemical interactions, without preventing activation of oxidative stress-responsive pathways.

#### 2.5.2 Autophagy

Inhibition of carbonyl stress and AGE-associated accumulation emerged as one of the biological outcomes of Ge-132 exposure [59,60,62]. Previous literature suggested improved cellular resilience and reduced stress-induced damage across several *in-vitro* models, including dermal fibroblasts and reproductive systems, indirectly pointing toward enhanced handling of damaged biomolecules [20,21,24,44,61,67,84]. The mechanism of autophagy is the natural process responsible for “cleaning” cell environment from toxic accumulation, and elimination of AGEs goes through this pathway [76].

To determine whether the modulation of glycative stress by Ge-132 can be associated with transcriptional changes in autophagy-related pathways, we examined genes involved in cargo recognition, pathway initiation, and autophagosome formation, as *SQSTM1, CALCOCO1, ULK1*, and *ATG-*related genes as representative of selective autophagy and early pathway activation (Figure 7b and S1).

MGO exposure induced a light but significant increase in *SQSTM1* and *CALCOCO1* expression. These genes encode cargo-recognition receptors that label damaged proteins and organelles for degradation, indicating activation of selective autophagy-related transcriptional responses under glycative stress. *SQSTM1* expression was further increased in the Ge-132+MGO condition, while *CALCOCO1* remained at the same level as MGO alone. The further increase in *SQSTM1* expression under combined Ge-132 and MGO exposure may indicate reinforcement of cargo-recognition responses under glycative stress.

At the level of pathway initiation, *ULK1* and *ATG4* were upregulated under MGO exposure. *ULK1* regulates autophagy initiation, and *ATG4* is involved in LC3 processing, indicating activation of early autophagy signaling. In the presence of Ge-132 the magnitude of this induction was not significantly reduced.

Genes associated with autophagosome elongation and membrane expansion (*ATG2B, ATG3, ATG5, and ATG7)* showed no significant variation across experimental conditions, suggesting that transcriptional activation is mainly restricted to early stages of the autophagy pathway (Supplemental Figure S1).

Overall, these results indicate that MGO-induced glycative stress is associated with selective transcriptional activation of early autophagy-related components, predominantly at the level of cargo recognition and pathway initiation. Within this context, Ge-132 may exert a context-dependent modulatory effect. In particular, we observe that Ge-132 does not substantially alter the transcriptional activation induced by MGO.

#### 2.5.3 Glyoxalase system and lysosomal pathway

To further evaluate detoxification and degradation capacity under glycative stress and the effect of Ge-132 on these processes, we analyzed the expression of key components of the glyoxalase system and lysosomal-associated genes, as GLO1, GLO2, and DJ-1 (Figure 8a).

**Figure 8.**
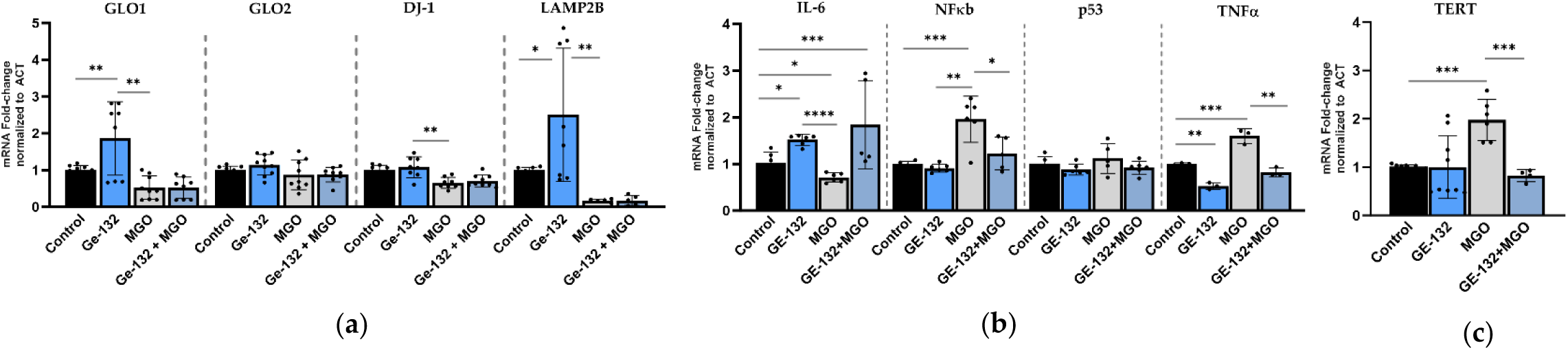
Transcriptional modulation of detoxification, lysosomal, and inflammatory pathways. (**a**) Expression of glyoxalase (*GLO1, GLO2, DJ-1)* and lysosomal-associated gene *LAMP2B*. Ge-132 increased GLO1 and LAMP2B expression from their basal conditions, while MGO exposure reduced *GLO1, DJ-1*, and *LAMP2B* levels. This reduction was maintained in the Ge-132+MGO condition. *GLO2* remained at the basal level across all conditions. (**b**) Expression of inflammatory markers (*IL-6, NF-κB, p53, TNFα*). MGO exposure increased *NF-κB* and *TNFα* expression, while reducing *IL-6* levels. Under combined Ge-132+MGO treatment, *NF-κB* and *TNFα* expression decreased toward control levels, whereas IL-6 expression was significantly increased compared to control. (**c**) Expression of TERT. MGO exposure increased TERT expression, whereas combined Ge-132+MGO treatment significantly reduced its expression compared to MGO alone. Data are normalized to actin and presented as mean of three biological replicates ± SD. Student’s t-test was used for statistical significance comparing each individual treatment *vs* related control, significance is indicated.

*GLO1* expression increased under Ge-132 treatment relative to basal conditions. GLO1 is the main enzyme responsible for methylglyoxal detoxification and that limits the formation of AGEs [85,86]. The induction of this gene under Ge-132 effect suggests a potential increase in detoxification capacity at the transcriptional level. In contrast, MGO exposure resulted in a marked downregulation of *GLO1* that is not countered by pretreatment with Ge-132. *GLO2* expression remained at the control level, across all experimental conditions.

*DJ-1* expression significantly decreased under MGO exposure and was not counteracted by the presence of Ge-132. DJ-1 functions as a redox-sensitive protein involved in both antioxidant defense and the detoxification of reactive carbonyl species [87,88]. Its persistent downregulation under MGO exposure, even in presence of Ge-132, points toward the idea that the reduction in AGEs accumulation is not primarily mediated by restoration of intracellular detoxification pathways but rather by upstream biochemical mechanisms.

Among lysosomal genes, *LAMP2B* expression significantly increased in response to Ge-132 alone. *LAMP2B* is associated with lysosomal function and autophagy-related degradation processes, suggesting enhanced basal lysosomal activity [89]. However, MGO exposure induced a pronounced reduction in *LAMP2B* expression, which persisted in the Ge-132+MGO condition. *LAMP2A* and *2C* showed a non-significant change of expression across the treatments (Supplemental Figure S1).

Overall, these results indicate that glycative stress is associated with reduced expression of key detoxification and lysosomal components, while Ge-132 primarily affects these pathways under non-stressed conditions.

#### 2.5.4 Inflammatory response

Animal studies have reported immunomodulatory properties of organogermanium compounds, including suppression of inflammasome activation, modulation of antiviral sensing pathways [65] and enhanced macrophage-mediated phagocytic activity [24]. Additional *in-vivo* work demonstrated a reduction of inflammatory damage in disease models [58]. To further assess whether glycative stress and its modulation by Ge-132 may influence inflammatory signaling, we analyzed the expression of key genes involved in pro-inflammatory pathways [82,90–94]. We found that glycative stress also altered inflammatory signaling in a selective and non-coordinated manner. For instance, MGO exposure increased *NF-κB* and *TNFα* expression together with *TERT*, indicating activation of stress-associated signaling pathways (Figure 8). The lack of *p53* activation, indicates only engagement of stress-related inflammatory signaling. In contrast, *IL-6* expression decreased under MGO exposure but showed partial recovery in the presence of Ge-132. These data suggest that glycative stress does not elicit a canonical pro-inflammatory response but rather a selective and asymmetric inflammatory profile. We observe activation of upstream inflammatory signaling without full downstream coordination, suggesting a non-uniform inflammatory response. However, a cell-type specific response cannot be excluded, so that the findings may not be generalizable across cell types.

Ge-132 alone did not induce a canonical pro-inflammatory response. Instead, a selective increase in *IL-6* expression was observed after cell treatment. However, no transcriptional changes are detected in canonical inflammatory mediators such as *TNFα, NF-κB,* or *p53*. MGO treatment induces a glycative stress and induces *NF-κB* and *TNFα* expression, but not *IL-6* or *p53*, still in line with a no clear activation of the pro-inflammatory pathway. Under combined Ge-132 and MGO exposure, the expression of *NF-κB* and *TNFα* was attenuated compared to MGO alone, while *IL-6* levels increased above control levels.

Overall, these data support a model in which glycative stress induces a selective and partially uncoupled inflammatory response, and Ge-132 acts as a context-dependent modulator that attenuates stress-induced signaling without restoring a coordinated inflammatory program.

## 3. Discussion

Several reports (Table 1 and Supplemental Table 1) have described redox and anti-inflammatory effects of Ge-132 across heterogeneous biological systems. Nevertheless, the field still lacks a unified mechanistic framework under standardized conditions. In the present study, we re-evaluated these proposed molecular targets within a single cellular model under controlled glycative stress, revealing a context-dependent modulation rather than uniform pathway activation.

For instance, the integration of biochemical and transcriptional analyses supports a model in which glycative stress may represent a functional bottleneck in cellular stress-response pathways. Our data confirm that exposure to methylglyoxal (MGO) promotes the accumulation of AGEs-modified proteins (Figure 6b) while simultaneously impairing key detoxification systems, including the glyoxalase components *GLO1* and *DJ-1*, together with lysosomal-associated machinery such as *LAMP2B* (Figure 8a). Thus, glycative stress not only increases the burden of damaged biomolecules but also compromises the systems required for their clearance.

In parallel, glycative stress triggers selective activation of autophagy-related pathways, mainly at the level of cargo recognition and early pathway initiation (Figure 7). The upregulation of *SQSTM1* and *CALCOCO1*, together with increased expression of upstream regulators such as *ULK1* and *ATG4*, indicates activation of adaptive stress responses. However, this is not accompanied by organized transcriptional activation of the core machinery required for autophagosome elongation and maturation (Supplemental Figure S1). Combined with reduced lysosomal capacity, this transcriptional pattern reveals only a selective transcriptional activation of early autophagy-related components.

Glycative stress also alters inflammatory signaling. MGO exposure increased *NF-κB*, *TNFα*, and *TERT* expression, indicating activation of stress-related and adaptive signaling pathways, while IL-6 expression was reduced (Figure 8 b,c). The increase in *TERT*, together with the absence of p53 activation, further supports a shift toward stress adaptation or survival-associated signaling. This pattern may be consistent with a non-coordinated or incomplete stress response. In this context, it is noteworthy that the transcriptional profiles observed in our model suggest a differential balance between inflammatory and antioxidant pathways. Under glycative stress conditions, *NF-κB* expression increased, whereas *NRF2* induction remained relatively limited. In contrast, under combined Ge-132 and MGO exposure, *NF-κB* expression was attenuated while *NRF2* expression showed a modest relative increase. Importantly, given that NRF2 activity is primarily regulated at the post-transcriptional level (KEAP1 system), these observations should be interpreted as changes in *NRF2*-related transcription rather than direct evidence of pathway activation. This pattern may be coherent with previously described interactions between *NF-κB* and *NRF2* signaling in which the NF-κB subunit p65 can modulate *NFE2L2* transcription in a context-dependent manner [95]. Although this interaction was not directly assessed in the present study, it provides a possible framework to interpret the partial and non-coordinated activation of stress-response pathways observed under these conditions. This shift in signaling balance under combined Ge-132 and glycative stress conditions, may contribute to the observed dissociation between biochemical damage reduction and sustained activation of stress-response pathways.

Within this framework, Ge-132 appears to exert a context-dependent modulatory effect, with features consistent with a priming-like effect under basal conditions. Under basal conditions, Ge-132 enhances the expression of selected detoxification and lysosomal components, including *GLO1* and *LAMP2B*, suggesting a potential increase in intrinsic capacity to respond to subsequent stress. Under glycative conditions, Ge-132 significantly reduces intracellular AGEs accumulation at the protein level. However, this reduction in glycative burden is not accompanied by transcriptional restoration of key detoxification and degradation systems, as *GLO1, DJ-1*, and *LAMP2B* remain transcriptionally suppressed under MGO exposure.

Notably, the persistence of *NRF2*-related transcriptional activity despite reduced AGE accumulation further supports a dissociation between biochemical damage reduction and cellular stress signaling.

In addition to alterations in detoxification and autophagy, glycative stress also affected inflammatory signaling in a selective and non-synchronized manner. MGO exposure increased *NF-κB* and *TNFα* expression, indicating activation of stress-related inflammatory pathways, while *IL-6* expression was reduced (Figure 8b). This divergent pattern suggests that glycative stress does not trigger a canonical pro-inflammatory response but rather an asymmetric activation of specific signaling components, consistent with a constrained and functionally incomplete stress response. In this context, Ge-132 fine-tunes inflammatory signaling rather than suppressing it, in a context-dependent manner.

Importantly, Ge-132 does not revert the transcriptional regulation imposed by MGO in autophagy detoxification and lysosomal pathways, suggesting that glycative stress impairs cellular homeostasis but Ge-132 does not prevent these alterations nor restore the transcriptional profile of these pathways under glycative stress (Figure 7 and 8).

Rather than restoring canonical cellular homeostasis, we propose that Ge-132 could partially rebalance the system at the level of biochemical damage, while cellular stress-response pathways remain transcriptionally active. These findings should be interpreted within the scope of the present study, which was designed to examine biochemical and transcriptional responses in a standardized cellular model, rather than provide an exhaustive and functional characterization of all downstream pathways.

Overall, our findings support a model in which glycative stress generates a restricted cellular state characterized by accumulation of damaged biomolecules, incomplete activation of autophagy at the transcriptional level, suppression of detoxification and lysosomal systems, and dysregulated inflammatory signaling (Figure 9). Within this context, Ge-132 reduces glycative damage without restoring cellular processing capacity, supporting a functional uncoupling between biochemical damage reduction and coordinated stress-response recovery.

**Figure 9.**
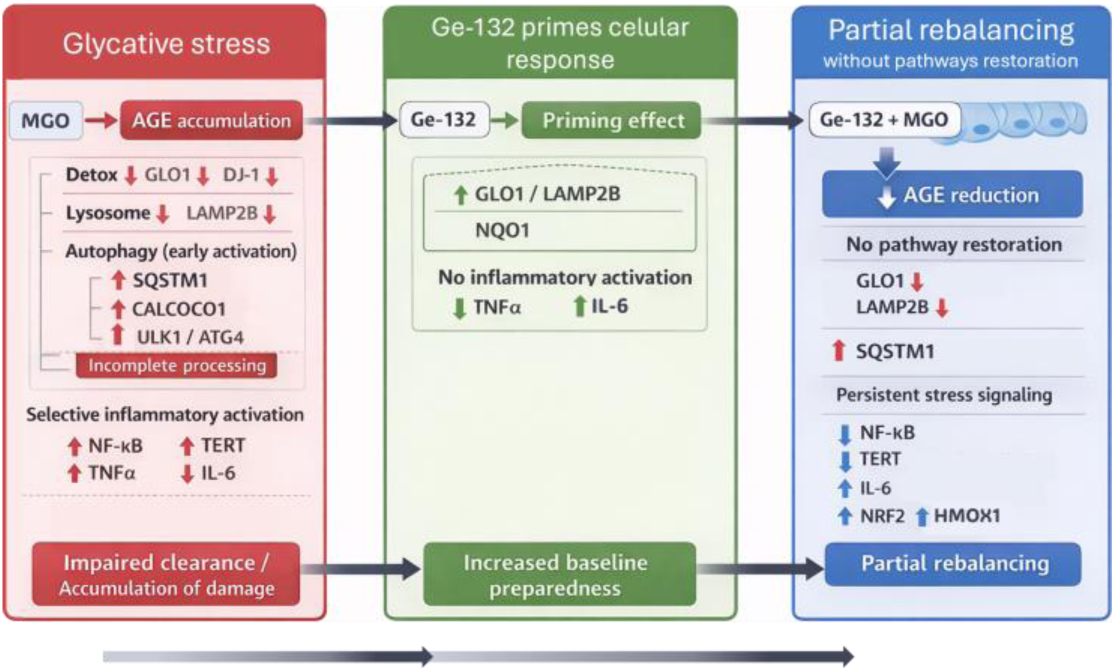
Proposed mechanistic model of Ge-132 activity under glycative stress. Exposure to methylglyoxal (MGO) induces accumulation of AGEs and represents a functional bottleneck in cellular stress-response pathways, suppressing key detoxification systems (*GLO1, DJ-1*) and lysosomal components (*LAMP2B*), while triggering selective activation of early autophagy-related processes (*SQSTM1, CALCOCO1, ULK1, ATG4*) without corresponding activation of downstream autophagy machinery. In parallel, inflammatory signaling is partially activated, as indicated by increased *NF-κB, TNFα* and *TERT* expression, while *IL-6* levels are reduced, coherent with a non-canonical inflammatory response. Under Ge-132 conditions selected detoxification and lysosomal genes are induced, hypothesizing a priming-like effect on cellular defense capacity. Under glycative stress, in cell pretreated with Ge-132, intracellular AGEs accumulation is reduced, and autophagy- and inflammation-related signaling remains transcriptionally active but not fully coordinated. Overall, Ge-132 does not restore the transcriptional profiles of detoxification and lysosomal pathways under glycative stress.

Beyond its descriptive value, the observed uncoupling between reduced glycative damage and sustained stress signaling may have biological relevance. Maintaining stress-response pathways active, even when damage is reduced, could help cells remain prepared to cope with ongoing or fluctuating stress, whereas an early shutdown of these responses might limit adaptation to subsequent challenges. In this context, the persistence of stress-related gene expression may reflect a protective, although not fully coordinated, response.

While our data do not directly demonstrate a causal uncoupling mechanism, they support a dissociation at the level of biochemical damage and transcriptional response under the conditions tested. Further validation at the protein level and through functional assays will be required to confirm the activity and integration of the pathways identified. In addition, the present findings are limited to a single, albeit consistent, cellular model. However, the use of a single fibroblast model limits extrapolation to other cell types, particularly those with distinct metabolic or inflammatory profiles. Nevertheless, the results support the idea that cellular stress responses are dynamic and do not necessarily resolve in parallel with biochemical changes.

## 4. Materials and Methods

### 4.1 Systematic literature review: search and selection

Article search and selection were conducted in accordance with PRISMA guidelines [96]. The literature search was designed as a multi-step evidence retrieval and screening strategy to identify original studies on organic germanium compounds, particularly Ge-132, in relation to biologically relevant effects associated with human health. Searches were performed across Web of Science, Scopus, PubMed, Europe PMC, and additional RIS-exported records from indexed databases, including ScienceDirect- and Embase-derived libraries, covering the last 15 years (2011–2025/2026, depending on database availability at the time of retrieval).

Two complementary search approaches were applied. A more restrictive search focused on human-related and clinically relevant studies using combinations of the terms germanium, organic germanium, Ge-132, carboxyethyl germanium, germanium sesquioxide, and carboxyethyl germanium sesquioxide in title/abstract fields together with human- or patient-related terms. A broader search incorporated biologically relevant contexts, including oxidative stress, antioxidant activity, inflammation, aging, autophagy, glycative stress, therapy, and pharmacological effects, while retaining Ge-132-related compound terms. Searches were limited to original articles and, where possible, filtered by biomedical and health-related research areas.

Record deduplication, and preliminary filtering were assisted by Julius AI, whereas final study selection was based on manual review according to predefined eligibility criteria. Because the aim of the review was to map both mechanistic and translational evidence, the screening process retained not only human studies but also relevant in-vitro and in-vivo experimental studies addressing the biological effects of Ge-132 within the predefined scope. Studies were considered eligible when the title, abstract, or keywords included at least one Ge-132-related term together with a relevant biological context, such as human/patient, clinical, in-vitro, in-vivo, mouse/mice, or rat-related terms.

Eligibility was restricted to original research articles published within the last 15 years that investigated organic germanium compounds, particularly Ge-132 or closely related denominations, and addressed outcomes relevant to human health or mechanistic biomedical interpretation, including oxidative stress, glycative/carbonyl stress, inflammation, autophagy, aging, pharmacological effects, or related cellular responses. Non-original publications, studies focused exclusively on inorganic germanium, non-biomedical applications, agricultural or plant systems, and false-positive records in which “Ge-132” referred to a non-compound identifier were excluded. Duplicate records were removed by DOI-based deduplication followed, when necessary, by title-based normalization. Retained studies were used for qualitative evidence mapping and data extraction, including publication of metadata, experimental model, sample type, dosage, and key biological outcomes.

Given the heterogeneity of the included literature, the review question was framed using an adapted PICO approach [97] focused on the biological effects of organic germanium compounds, particularly Ge-132, across human, animal, and in-vitro models, with emphasis on outcomes related to oxidative stress, glycative/carbonyl stress, inflammatory signaling, autophagy, aging, and associated molecular or physiological responses.

#### 4.1.1 Risk of bias assessment (RoB)

RoB was assessed using an adapted version of the Critical Appraisal Skills Programme (CASP) checklist, modified to fit the experimental nature of the included studies [70,98]. Data extraction were managed using the Covidence systematic review software [99]. After full-text screening and included the following domains: sequence generation, allocation concealment, incomplete outcome data, selective outcome reporting, and other sources of bias. Risk judgments were classified as low, high, or unclear. Because the included studies comprised heterogeneous experimental settings, risk-of-bias assessment was conducted separately for *in-vitro* and *in-vivo* studies in order to preserve comparability within each evidence stream, and to support structured evidence mapping. The domain “other sources of bias” was used to capture potential concerns related to structural or contextual factors not fully addressed by the core methodological domains, including limited independent replication, compound sourcing, and possible funding- or collaboration-related influences.

### 4.2 Cell culture conditions

Cells were cultured in Dulbecco’s Modified Eagle Medium (DMEM, GIBCO) supplemented with 10% fetal bovine serum (FBS), 1% penicillin-streptomycin mix (10,000 U/mL), 1% MEM non-essential amino acids (GIBCO), and 1% sodium pyruvate 100 mM (GIBCO), and maintained at 37 °C in a humidified atmosphere containing 5% CO_2_. When required by the treatment, the medium was supplemented with Ge-132 at the concentration indicated in each case (Glentham Life Sciences, Ref. 7W-GP2575). Ge-132 powder was dissolved in DMEM, the pH was equilibrated to 6.8, and the solution was filtered and used freshly. Methylglyoxal solution (MGO, Sigma-Aldrich) was diluted to 2mM final concentration in DMEM and filter sterilized before apply the treatment to the cells.

### 4.3 Analysis of Cytotoxicity

Mouse embryonic fibroblasts (MEFs), Human Lens Epithelial Cells (HLECs), and human Retinal Pigment Epithelial cells (ARPE19), were kindly provided by E. Bejarano’s group [100]. Cells were generally seeded in 96-well plates at a density of 5 × 10³ cells per well and incubated for 24 h. Cells were then exposed to different concentrations of Ge-132, ranging from 0 to 20 mM, for 24 h. After treatment, cell proliferation and viability were assessed using the 3-(4,5-dimethylthiazol-2-yl)-2,5-diphenyl tetrazolium bromide (MTT) assay. Briefly, 5 μL of MTT solution (20 mg/mL) was added to each well, and the plate was incubated at 37 °C for 2 h. Then, 200 μL of dimethyl sulfoxide was added to solubilize the formazan crystals, and the plate was maintained in the dark at room temperature for 30 min. Absorbance was measured at 570 nm using a Varioskan LUX plate reader (Thermo Fisher Scientific, Vantaa, Finland). Data were collected and analyzed normalizing to control group. Each assay was repeated in triplicate.

### 4.4 Quantitative Real-Time PCR Assays

MEFs were grown in Petri dishes until reaching approximately 80% confluence. Then, Ge-132 treatment was supplied at 5 mM, for 24 hr. MGO was prepared at 2mM and added to the cells for 2 hours. These conditions are consistent with previously reported acute glycative stress models [76;103]. After treatments, cells were washed once with cold 1× PBS and scraped in the same buffer. After gentle centrifugation, the medium was removed and the cell pellets were stored at −80 °C until processing. Total RNA was isolated using the NZY Total RNA Kit (NZYTech, Lisboa, Portugal), according to the manufacturer’s instructions. Subsequently, 1 μg of total RNA from each sample was reverse-transcribed using the NZY Reverse Transcriptase Kit (NZYTech, Lisboa, Portugal). Quantitative real-time PCR (qRT-PCR) was performed to assess the mRNA expression levels of the genes reported in Supplemental Table S2. Reactions were carried out using NZY Speedy qPCR Green Master Mix (NZYTech, Lisboa, Portugal) in a QuantStudio™ 5 Real-Time PCR System. Relative gene expression was calculated after internal normalization to the housekeeping gene β-actin and referred to the control condition.

### 4.5 In-vitro antiglycation assay

The antiglycation activity of Ge-132 was evaluated using a cell-free BSA–glucose model based on a previously described fluorescence assay, with minor adaptations [101]. Briefly, bovine serum albumin (BSA, 10 mg/mL) and anhydrous glucose (90 mg/mL) were prepared separately in phosphate-buffered saline (PBS, pH 7.4). Reaction mixtures were assembled by combining BSA, glucose, and Ge-132 at the concentrations indicated in the corresponding figure. Aminoguanidine (AG) was included as a positive control for inhibition of AGE formation, whereas control reactions were prepared in parallel in the absence of Ge-132. Sodium azide (0.01%) was added to prevent microbial contamination. The reaction mixtures were incubated at 37 °C for 7 days in the dark.

After incubation, AGEs formation was quantified fluorometrically using excitation and emission wavelengths of 360 and 420 nm, respectively. Measurements were performed in triplicate. The percentage inhibition of AGE formation was calculated according to the formula: *Inhibition (%) = [(F0 – F1) / F0] × 100*, where *F0* represents the fluorescence intensity of the glycated control and *F1* the fluorescence intensity of the Ge-132-treated sample.

### 4.6 Protein purification, quantification and Western blot

Protein extracts were prepared in the presence of protease inhibitors (Roche, Complete tablets Ref. 04693132001) in a PBS1x buffer and quantified using a colorimetric BCA protein assay (Pierce, BCA Protein Assay Kit, REF 23225), according to the manufacturer’s instructions. Samples and standards were analyzed in duplicate, and absorbance values were used to calculate protein concentration from a calibration curve. Protein quantification was performed to normalize sample loading for subsequent electrophoretic analysis.

Protein expression and specifically MG-modification, was analyzed by Western blot. Equal amounts of total protein were mixed with denaturing Laemmli SDS buffer, heat-denatured, and separated by SDS-PAGE. Proteins were then transferred onto a nitrocellulose-membrane, which was stained with Ponceau solution to verify transfer efficiency and equal loading, blocked with TBS-T-milk 5% solution, and incubated with anti-MG primary antibody (Mouse Anti-Methylglyoxal Monoclonal Antibody, Cell Biolabs, ref. STA-011), at 1:1000 dilution in TBST-milk 5% buffer, overnight at 4 °C. After washing, membranes were incubated with the secondary antibody at 1:20000 dilution, for 2 hours at room temperature. Immunodetection was realized with ECL Prime Detection kit (Amersham), and visualized in a digital imaging system (Omega Lum C). Bands intensity were analyzed densitometrically with ImageJ (Fiji) software [102].

### 4.7 Statistical analysis and software

Statistical analyses were performed using GraphPad Prism (version 10.1.2, GraphPad Software, LLC). Data are presented as mean ± standard deviation (SD) from at least three independent experiments. Outliers in the technical replicates were identified by ROUT method, applying a Q=0.5%.

For comparisons involving more than two groups, statistical significance was assessed using one-way analysis of variance (ANOVA) followed by Dunnett’s multiple comparison test to compare each treatment condition with the corresponding control.

For experiments involving only two groups, unpaired two-tailed Student’s t-test was used. Data distribution was assumed to be approximately normal based on experimental design and sample size. A p-value < 0.05 was considered statistically significant and the corresponding data are marked by an asterisk following this correspondence: 0.033(*), 0.002(**), 0.002(***), 0.0001 (****).

## 5. Conclusions

In conclusion, the present study provides an integrated analysis combining literature mapping and experimental validation to examine the biological effects of Ge-132 under glycative stress conditions. Our results confirm that Ge-132 reduces the accumulation of glycative damage at the biochemical level. However, this effect is not accompanied by restoration of coordinated transcriptional responses across key pathways involved in detoxification, autophagy, and inflammation. Instead, we observe a selective and context-dependent modulation of gene expression, suggesting that cellular stress-response pathways remain active despite reduced damage.

These findings support a dissociation between glycative damage reduction and cellular regulatory responses under the conditions tested. While further validation at the protein and functional levels will be required to fully characterize the mechanisms involved, this work provides a conceptual framework that may help guide future studies aimed at integrating biochemical and cellular perspectives in the study of organogermanium compounds..

## Supplementary Materials

The following supporting information can be downloaded at: https://www.mdpi.com/article/doi/s1, Figure S1: Relative expression of autophagy and lysosomal-associated genes. Table S1: Ge-132 systematic review Full tables, Table S2: qRT-PCR Oligonucleotides

## Author Contributions

Conceptualization, A.L.; methodology, A.P., A.L.; validation, A.P., A.M., G.G. and Y.F.A.E; formal analysis, A.P., Y.F.A.E and A.L.; investigation, A.P., A.M., G.G. and Y.F.A.E.; data curation, A.L.; writing—original draft preparation, A.L., G.G. and Y.F.A.E; writing—review and editing, A.L., G.G. and Y.F.A.E.; visualization, G.G. and Y.F.A.E.; supervision, A.L.; project administration, A.L.; funding acquisition, A.L.. All authors have read and agreed to the published version of the manuscript.

## Data Availability Statement

The original contributions presented in this study are included in the article. Further inquiries can be directed to the corresponding author.

## Acknowledgments

The authors sincerely thank Dr. Teresa López de Coca Pérez for sharing her expertise and for her guidance in the appropriate selection and analysis of the articles included in the systematic review section of this manuscript. A special acknowledgement for Dr Eloy Bejarano who kindly provided all the cell lines used in this work, and for revising this manuscript. The authors also acknowledge the students of the Medicine Degree Program at CEU Cardenal Herrera University (CEU-UCH) who enthusiastically participated in this study within a university training context, although they did not directly contribute to the generation of the reported results. During the preparation of this manuscript, the authors used Oreate AI to assist in Graphical abstract, Figures 1 and Figure 9 design; Elicit AI has been used to assist in table organization; Julius AI assisted literature deduplication after manual search, and preliminary filtering applying inclusion criteria. The authors have reviewed and edited the output and take full responsibility for the content of this publication.

## Appendix A

### Appendix A.1

**Table A1. Ge-132 systematic review Full tables**

**Table A2. qRT-PCR Oligonucleotides**

---

**Figure S1.**
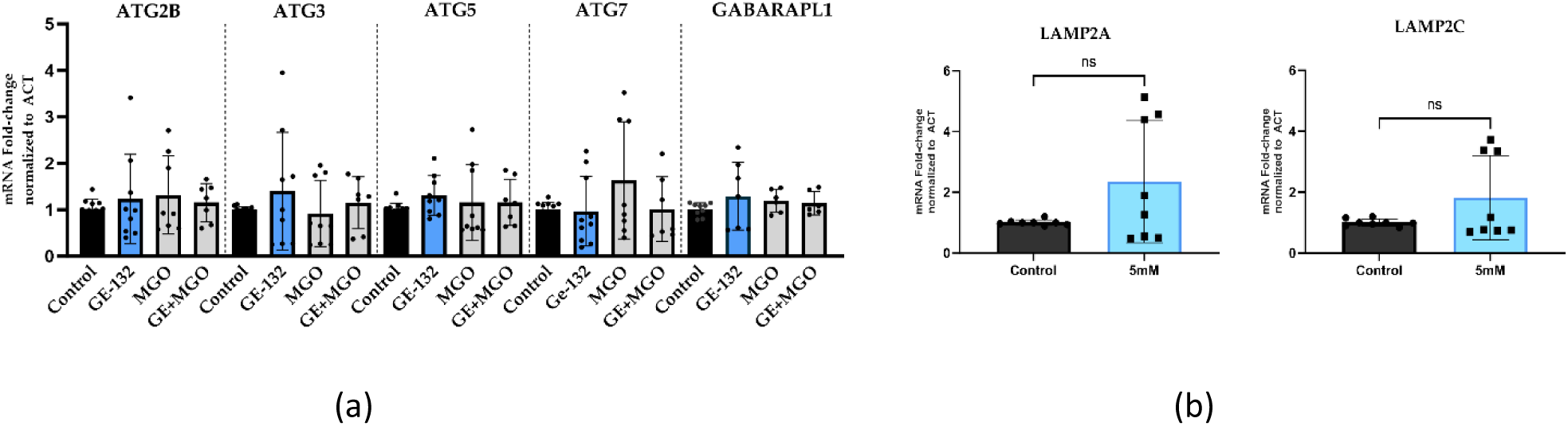
Relative expression of autophagy and lysosomal-associated genes. (a) MGO induced upregulation of an early regulatory components of autophagy (*ATG4*), whereas genes involved in autophagosome elongation (i.e. *ATG2B, ATG3, ATG5, ATG7*) showed limited or no consistent changes. GABARAPL1, which is mainly involved in autophagosomal membrane expansion, cargo enclosure, and later maturation/fusion steps, is not displaying any change across the treatments. (b) Expression of lysosomal-associated genes LAMP2A and 2C remained unaltered across Ge-132 and MGO treatments. Expression levels were normalized to Actin and are presented relative to control. Data represent mean ± SD from 3 independent experiments and an average of 6–10 technical replicates. Statistical significance indicated as *P < 0.05.

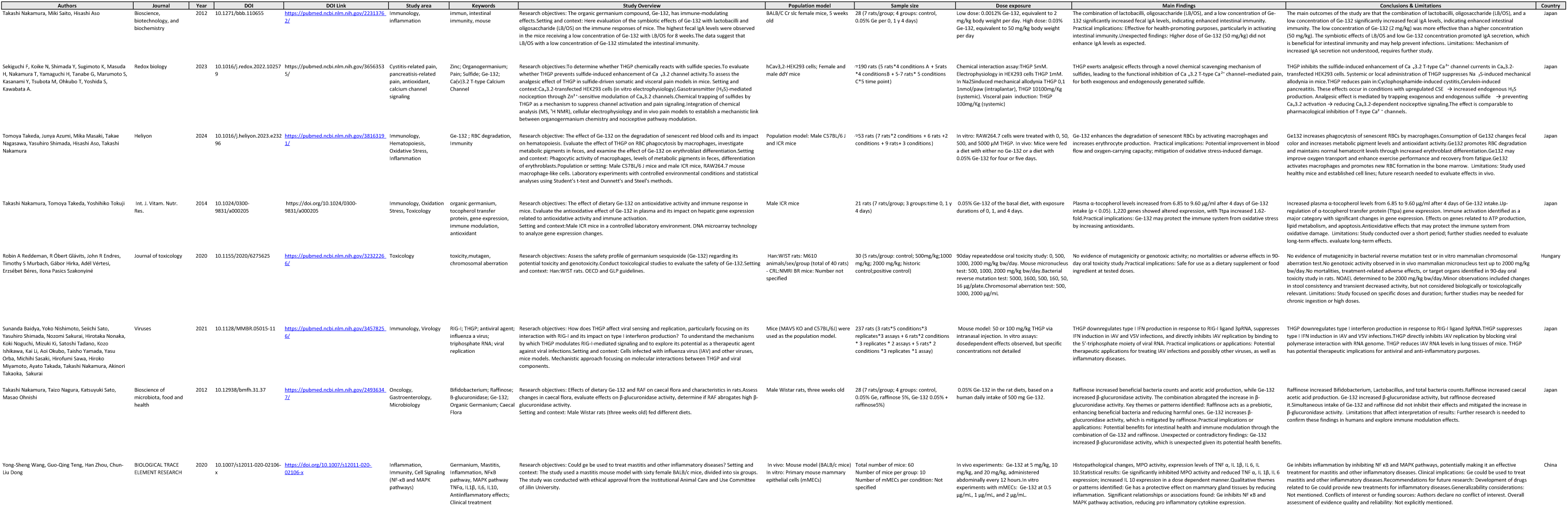
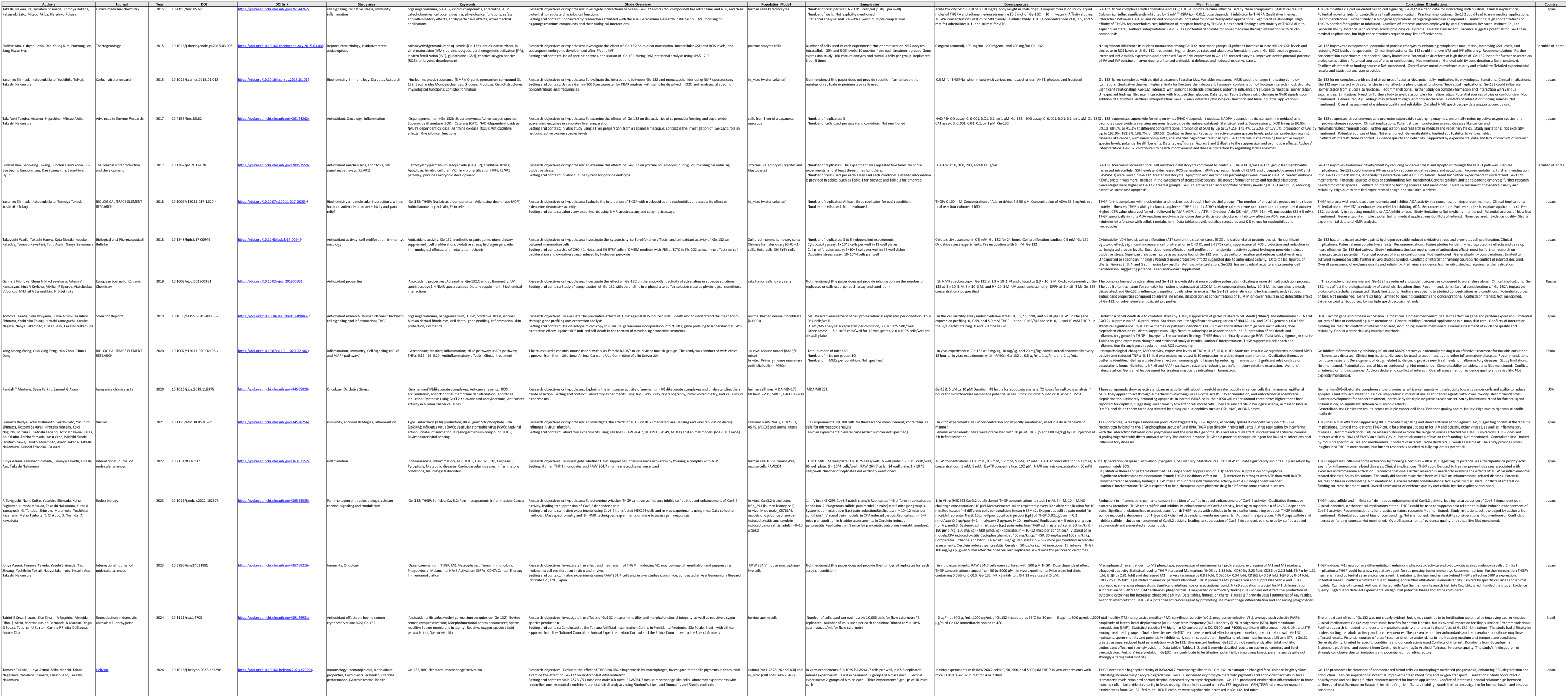

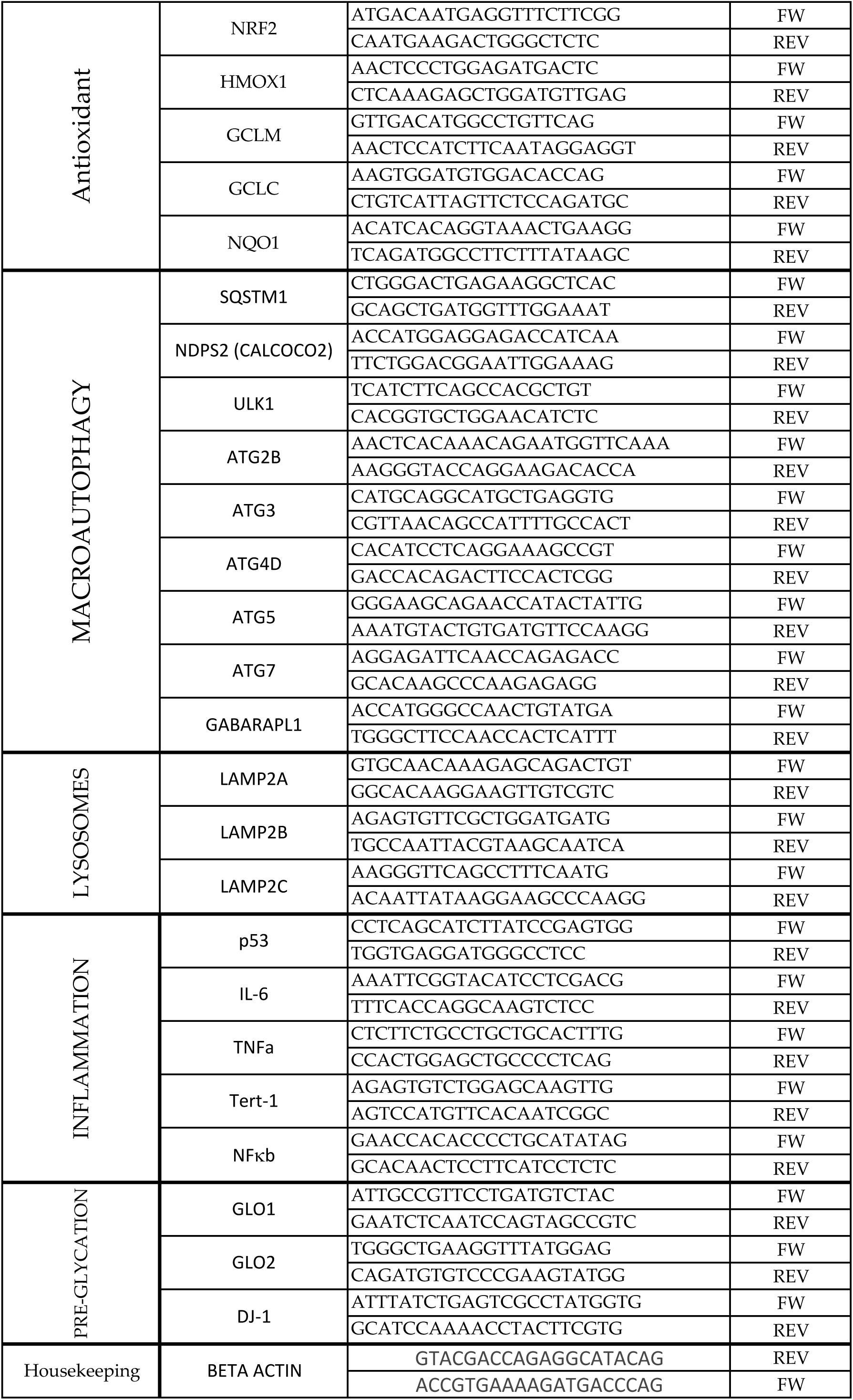

