## Supplemental Tables for "Organic Germanium (Ge-132) reduces glycative damage while maintaining cellular stress signaling: evidence of functional dissociation"

**Supplemental Figure S1**

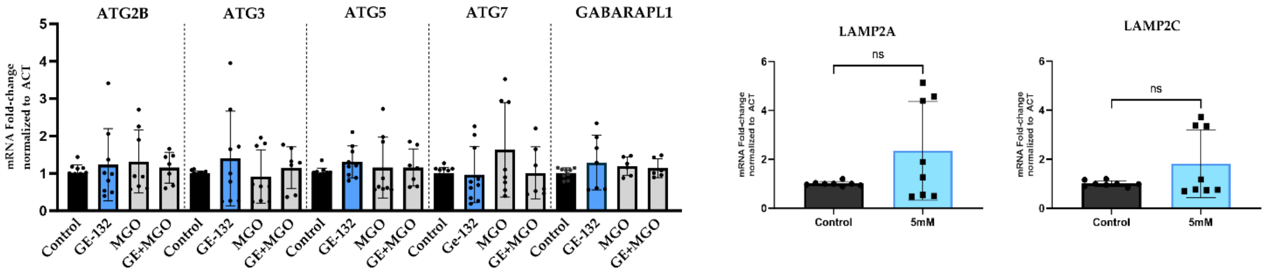


(a) (b)

Figure S1. Relative expression of autophagy and lysosomal-associated genes. (a) MGO induced upregulation of an early regulatory components of autophagy (*ATG4*), whereas genes involved in autophagosome elongation (i.e. *ATG2B, ATG3, ATG5, ATG7*) showed limited or no consistent changes. GABARAPL1, which is mainly involved in autophagosomal membrane expansion, cargo enclosure, and later maturation/fusion steps, is not displaying any change across the treatments. (b) Expression of lysosomal-associated genes LAMP2A and 2C remained unaltered across Ge-132 and MGO treatments.

Expression levels were normalized to Actin and are presented relative to control. Data represent mean ± SD from 3 independent experiments and an average of 6–10 technical replicates. Statistical significance indicated as $^*P < 0.05$.

| Authors | Journal | Year | DOI | DOI Link | Study area | Keywords | Study Overview | Population model | Sample size | Dose exposure | Main Findings | Conclusions & Limitations | Country |
|---|---|---|---|---|---|---|---|---|---|---|---|---|---|
| Takashi Nakamura, Miki Saito, Hisashi Aso | Bioscience, biotechnology, and biochemistry | 2012 | 10.1271/bbb.110655 | https://pubmed.ncbi.nlm.nih.gov/22313762/ | Immunology, inflammation | immun, intestinal immunity, mouse | Research objectives: The organic germanium compound, Ge-132, has immune-modulating effects.Setting and context: Here evaluation of the symbiotic effects of Ge-132 with lactobacilli and oligosaccharide (LB/OS) on the immune responses of mice. The highest fecal IgA levels were observed in the mice receiving a low concentration of Ge-132 with LB/OS for 8 weeks.The data suggest that LB/OS with a low concentration of Ge-132 stimulated the intestinal immunity. | BALB/C Cr slc female mice, 5 weeks old | 28 (7 rats/group; 4 groups: control, 0.05% Ge per 0, 1 y 4 days) | Low dose: 0.0012% Ge-132, equivalent to 2 mg/kg body weight per day. High dose: 0.03% Ge-132, equivalent to 50 mg/kg body weight per day | The combination of lactobacilli, oligosaccharide (LB/OS), and a low concentration of Ge-132 significantly increased fecal IgA levels, indicating enhanced intestinal immunity. Practical implications: Effective for health-promoting purposes, particularly in activating intestinal immunity.Unexpected findings: Higher dose of Ge-132 (50 mg/kg) did not enhance IgA levels as expected. | The main outcomes of the study are that the combination of lactobacilli, oligosaccharide (LB/OS), and a low concentration of Ge-132 significantly increased fecal IgA levels, indicating enhanced intestinal immunity. The low concentration of Ge-132 (2 mg/kg) was more effective than a higher concentration (50 mg/kg). The symbiotic effects of LB/OS and low Ge-132 concentration promoted IgA secretion, which is beneficial for intestinal immunity and may help prevent infections. Limitations: Mechanism of increased IgA secretion not understood, requires further study. | Japan |
| Sekiguchi F, Koike N, Shimada Y, Sugimoto K, Masuda H, Nakamura T, Yamaguchi H, Tanabe G, Marumoto S, Kasanami Y, Tsubota M, Ohkubo T, Yoshida S, Kawabata A. | Redox biology | 2023 | 10.1016/j.redox.2022.102579 | https://pubmed.ncbi.nlm.nih.gov/36563535/ | Cystitis-related pain, pancreatisis-related pain, antioxidant, calcium channel signaling | Zinc; Organogermanium; Pain; Sulfide; Ge-132; Ca(v)3.2 T-type Calcium Channel | Research objectives:To determine whether THGP chemically reacts with sulfide species.To evaluate whether THGP prevents sulfide-induced enhancement of $Ca_v3.2$ channel activity.To assess the analgesic effect of THGP in sulfide-driven somatic and visceral pain models in mice. Setting and context:$Ca_v3.2$-transfected HEK293 cells (in vitro electrophysiology).Gasotransmitter ($H_2S$)-mediated nociception through $Zn^{2+}$-sensitive modulation of $Ca_v3.2$ channels.Chemical trapping of sulfides by THGP as a mechanism to suppress channel activation and pain signaling.Integration of chemical analysis (MS, $^1H$ NMR), cellular electrophysiology and in vivo pain models to establish a mechanistic link between organogermanium chemistry and nociceptive pathway modulation. | hCav3,2-HEK293 cells; Female and male ddY mice | ≈190 rats (5 rats*4 conditions A + 5rats *4 conditionsB + 5-7 rats* 5 conditions C*5 time point) | Chemical interaction assay:THGP 5mM. Electrophysiology in HEK293 cells THGP 1mM. In Na2Sinduced mechanical allodynia THGP 0,1 1nmol/paw (intraplantar), THGP 10100mg/Kg (systemic). Visceral pain induction: THGP 100mg/Kg (systemic) | THGP exerts analgesic effects through a novel chemical scavenging mechanism of sulfides, leading to the functional inhibition of $Ca_v3.2$ T-type $Ca^{2+}$ channel–mediated pain, for both exogenous and endogenously generated sulfide. | THGP inhibits the sulfide-induced enhancement of $Ca_v3.2$ T-type $Ca^{2+}$ channel currents in $Ca_v3.2$-transfected HEK293 cells. Systemic or local administration of THGP suppresses $Na_2S$-induced mechanical allodynia in mice.THGP reduces pain in:Cyclophosphamide-induced cystitis,Cerulein-induced pancreatitis. These effects occur in conditions with upregulated CSE → increased endogenous $H_2S$ production. Analgesic effect is mediated by trapping exogenous and endogenous sulfide → preventing $Ca_v3.2$ activation → reducing $Ca_v3.2$-dependent nociceptive signaling.The effect is comparable to pharmacological inhibition of T-type $Ca^{2+}$ channels. | Japan |
| Tomoya Takeda, Junya Azumi, Mika Masaki, Takae Nagasawa, Yasuhiro Shimada, Hisashi Aso, Takashi Nakamura | Heliyon | 2024 | 10.1016/j.heliyon.2023.e23296 | https://pubmed.ncbi.nlm.nih.gov/38163191/ | Immunology, Hematopoiesis, Oxidative Stress, Inflammation | Ge-132 ; RBC degradation, Immunity | Research objective: The effect of Ge-132 on the degradation of senescent red blood cells and its impact on hematopoiesis. Evaluate the effect of THGP on RBC phagocytosis by macrophages, investigate metabolic pigments in feces, and examine the effect of Ge-132 on erythroblast differentiation.Setting and context: Phagocytic activity of macrophages, levels of metabolic pigments in feces, differentiation of erythroblasts.Population or setting: Male C57BL/6 J mice and male ICR mice, RAW264.7 mouse macrophage-like cells. Laboratory experiments with controlled environmental conditions and statistical analyses using Student's t-test and Dunnett's and Steel's methods. | Population model: Male C57BL/6 J and ICR mice | ≈53 rats (7 rats*2 conditions + 6 rats +2 conditions + 9 rats+ 3 conditions) | In vitro: RAW264.7 cells were treated with 0, 50, 500, and 5000 μM THGP. In vivo: Mice were fed a diet with either no Ge-132 or a diet with 0.05% Ge-132 for four or five days. | Ge-132 enhances the degradation of senescent RBCs by activating macrophages and increases erythrocyte production. Practical implications: Potential improvement in blood flow and oxygen-carrying capacity; mitigation of oxidative stress-induced damage. | Ge132 increases phagocytosis of senescent RBCs by macrophages.Consumption of Ge132 changes fecal color and increases metabolic pigment levels and antioxidant activity.Ge132 promotes RBC degradation and maintains normal hematocrit levels through increased erythroblast differentiation.Ge132 may improve oxygen transport and enhance exercise performance and recovery from fatigue.Ge132 activates macrophages and promotes new RBC formation in the bone marrow. Limitations: Study used healthy mice and established cell lines; future research needed to evaluate effects in vivo. | Japan |
| Takashi Nakamura, Tomoya Takeda, Yoshihiko Tokuji | Int. J. Vitam. Nutr. Res. | 2014 | 10.1024/0300-9831/a000205 | https://doi.org/10.1024/0300-9831/a000205 | Immunology, Oxidation Stress, Toxicology | organic germanium, tocopherol transfer protein, gene expression, immune modulation, antioxidant | Research objectives: The effect of dietary Ge-132 on antioxidative activity and immune response in mice. Evaluate the antioxidative effect of Ge-132 in plasma and its impact on hepatic gene expression related to antioxidative activity and immune activation.<br>Setting and context:Male ICR mice in a controlled laboratory environment. DNA microarray technology to analyze gene expression changes. | Male ICR mice | 21 rats (7 rats/group; 3 groups:time 0, 1 y 4 days) | 0.05% Ge-132 of the basal diet, with exposure durations of 0, 1, and 4 days. | Plasma α-tocopherol levels increased from 6.85 to 9.60 μg/ml after 4 days of Ge-132 intake ($p < 0.05$). 1,220 genes showed altered expression, with Ttpa increased 1.62-fold.Practical implications: Ge-132 may protect the immune system from oxidative stress by increasing antioxidants. | Increased plasma α-tocopherol levels from 6.85 to 9.60 μg/ml after 4 days of Ge-132 intake.Up-regulation of α-tocopherol transfer protein (Ttpa) gene expression. Immune activation identified as a major category with significant changes in gene expression. Effects on genes related to ATP production, lipid metabolism, and apoptosis.Antioxidative effects that may protect the immune system from oxidative damage. Limitations: Study conducted over a short period; further studies needed to evaluate long-term effects. evaluate long-term effects. | Japan |
| Robin A Reddeman, R Óbert Glávits, John R Endres, Timothy S Murbach, Gábor Hirka, Adél Vértesi, Erzsébet Béres, Ilona Pasics Szakonyiné | Journal of toxicology | 2020 | 10.1155/2020/6275625 | https://pubmed.ncbi.nlm.nih.gov/32322266/ | Toxicology | toxicity,mutagen, chromosomal aberration | Research objectives: Assess the safety profile of germanium sesquioxide (Ge-132) regarding its potential toxicity and genotoxicity.Conduct toxicological studies to evaluate the safety of Ge-132.Setting and context: Han:WIST rats. OECD and GLP guidelines. | Han:WIST rats: M610 animals/sex/group (total of 40 rats) - CRL:NMRI BR mice: Number not specified | 30 (5 rats/group: control; 500mg/kg;1000 mg/kg; 2000 mg/kg; historic control;positive control) | 90day repeateddose oral toxicity study: 0, 500, 1000, 2000 mg/kg bw/day. Mouse micronucleus test: 500, 1000, 2000 mg/kg bw/day.Bacterial reverse mutation test: 5000, 1600, 500, 160, 50, 16 μg/plate.Chromosomal aberration test: 500, 1000, 2000 μg/mL | No evidence of mutagenicity or genotoxic activity; no mortalities or adverse effects in 90-day oral toxicity study.Practical implications: Safe for use as a dietary supplement or food ingredient at tested doses. | No evidence of mutagenicity in bacterial reverse mutation test or in vitro mammalian chromosomal aberration test.No genotoxic activity observed in in vivo mammalian micronucleus test up to 2000 mg/kg bw/day.No mortalities, treatment-related adverse effects, or target organs identified in 90-day oral toxicity study in rats. NOAEL determined to be 2000 mg/kg bw/day.Minor observations included changes in stool consistency and transient decreased activity, but not considered biologically or toxicologically relevant. Limitations: Study focused on specific doses and duration; further studies may be needed for chronic ingestion or high doses. | Hungary |
| Sunanda Baidya, Yoko Nishimoto, Seiichi Sato, Yasuhiro Shimada, Nozomi Sakurai, Hirotaka Nonaka, Koki Noguchi, Mizuki Ki, Satoshi Tadano, Kozo Ishikawa, Kai Li, Aoi Okubo, Taisho Yamada, Yasu Orba, Michihi Sasaki, Hirofumi Sawa, Hiroko Miyamoto, Ayato Takada, Takashi Nakamura, Akinori Takaoka, Sakurai | Viruses | 2021 | 10.1128/MMBR.05015-11 | https://pubmed.ncbi.nlm.nih.gov/34578256/ | Immunology, Virology | RIG-I; THGP; antiviral agent; influenza a virus; triphosphate RNA; viral replication | Reserach objectives: How does THGP affect viral sensing and replication, particularly focusing on its interaction with RIG-I and its impact on type I interferon production? To understand the mechanisms by which THGP modulates RIG-I-mediated signaling and to explore its potential as a therapeutic agent against viral infections.Setting and context: Cells infected with influenza virus (IAV) and other viruses, mice models. Mechanistic approach focusing on molecular interactions between THGP and viral components. | Mice (MAVS KO and C57BL/6J) were used as the population model. | 237 rats (3 rats*5 conditions*3 replicates*3 assays + 6 rats*2 conditions * 3 replicates * 2 assays + 5 rats* 2 conditions *3 replicates *1 assay) | Mouse model: 50 or 100 mg/kg THGP via intranasal injection. In vitro assays: dosedependent effects observed, but specific concentrations not detailed | THGP downregulates type I IFN production in response to RIG-I ligand 3pRNA, suppresses IFN induction in IAV and VSV infections, and directly inhibits IAV replication by binding to the 5'-triphosphate moiety of viral RNA. Practical implications or applications: Potential therapeutic applications for treating IAV infections and possibly other viruses, as well as inflammatory diseases. | THGP downregulates type I interferon production in response to RIG-I ligand 3pRNA.THGP suppresses type I IFN induction in IAV and VSV infections.THGP directly inhibits IAV replication by blocking viral polymerase interaction with RNA genome. THGP reduces IAV RNA levels in lung tissues of mice. THGP has potential therapeutic implications for antiviral and anti-inflammatory purposes. | Japan |
| Takashi Nakamura, Taizo Nagura, Katsuyuki Sato, Masao Ohnishi | Bioscience of microbiota, food and health | 2012 | 10.12938/bmfh.31.37 | https://pubmed.ncbi.nlm.nih.gov/24936347/ | Oncology, Gastroenterology, Microbiology | Bifidobacterium; Raffinose; B-glucuronidase; Ge-132; Organic Germanium; Caecal Flora | Research objectives: Effects of dietary Ge-132 and RAF on caecal flora and characteristics in rats.Assess changes in caecal flora, evaluate effects on β-glucuronidase activity, determine if RAF abrogates high β-glucuronidase activity.<br>Setting and context: Male Wistar rats (three weeks old) fed different diets. | Male Wistar rats, three weeks old | 28 (7 rats/group; 4 groups: control, 0.05% Ge, raffinose 5%, Ge-132 0.05% + raffinose5%) | 0.05% Ge-132 in the rat diets, based on a human daily intake of 500 mg Ge-132. | Raffinose increased beneficial bacteria counts and acetic acid production, while Ge-132 increased β-glucuronidase activity. The combination abrogated the increase in β-glucuronidase activity. Key themes or patterns identified: Raffinose acts as a prebiotic, enhancing beneficial bacteria and reducing harmful ones. Ge-132 increases β-glucuronidase activity, which is mitigated by raffinose.Practical implications or applications: Potential benefits for intestinal health and immune modulation through the combination of Ge-132 and raffinose. Unexpected or contradictory findings: Ge-132 increased β-glucuronidase activity, which is unexpected given its potential health benefits. | Raffinose increased Bifidobacterium, Lactobacillus, and total bacteria counts.Raffinose increased caecal acetic acid production. Ge-132 increased β-glucuronidase activity, but raffinose decreased it.Simultaneous intake of Ge-132 and raffinose did not inhibit their effects and mitigated the increase in β-glucuronidase activity. Limitations that affect interpretation of results: Further research is needed to confirm these findings in humans and explore immune modulation effects. | Japan |
| Yong-Sheng Wang, Guo-Qing Teng, Han Zhou, Chun-Liu Dong | BIOLOGICAL TRACE ELEMENT RESEARCH | 2020 | 10.1007/s12011-020-02106-x | https://doi.org/10.1007/s12011-020-02106-x | Inflammation, Immunity, Cell Signaling (NF-κB and MAPK pathways) | Germanium, Mastitis, Inflammation, NFκB pathway, MAPK pathway TNFα, IL1β, IL6, IL10, Antiinflammatory effects; Clinical treatment | Research objectives: Could ge be used to treat mastitis and other inflammatory diseases? Setting and context: The study used a mastitis mouse model with sixty female BALB/c mice, divided into six groups. The study was conducted with ethical approval from the Institutional Animal Care and Use Committee of Jilin University. | In vivo: Mouse model (BALB/c mice)<br>In vitro: Primary mouse mammary epithelial cells (mMECs) | Total number of mice: 60<br>Number of mice per group: 10<br>Number of mMECs per condition: Not specified | In vivo experiments: Ge-132 at 5 mg/kg, 10 mg/kg, and 20 mg/kg, administered abdominally every 12 hours.In vitro experiments with mMECs: Ge-132 at 0.5 μg/mL, 1 μg/mL, and 2 μg/mL. | Histopathological changes, MPO activity, expression levels of TNF α, IL 1β, IL 6, IL 10.Statistical results: Ge significantly inhibited MPO activity and reduced TNF α, IL 1β, IL 6 expression; increased IL 10 expression in a dose dependent manner.Qualitative themes or patterns identified: Ge has a protective effect on mammary gland tissues by reducing inflammation. Significant relationships or associations found: Ge inhibits NF κB and MAPK pathway activation, reducing pro inflammatory cytokine expression. | Ge inhibits inflammation by inhibiting NF κB and MAPK pathways, potentially making it an effective treatment for mastitis and other inflammatory diseases. Clinical implications: Ge could be used to treat mastitis and other inflammatory diseases.Recommendations for future research: Development of drugs related to Ge could provide new treatments for inflammatory diseases.Generalizability considerations: Not mentioned. Conflicts of interest or funding sources: Authors declare no conflict of interest. Overall assessment of evidence quality and reliability: Not explicitly mentioned. | China |

| Authors | Journal | Year | DOI | DOI link | Study area | Keywords | Study Overview | Population Model | Sample size | Dose exposure | Main Findings | Conclusions & Limitations | Country |
|---|---|---|---|---|---|---|---|---|---|---|---|---|---|
| Takashi Nakamura, Yasuhiro Shimada, Tomoya Takeda, Katsuyuki Sato, Mitsuo Akiba, Haruhiko Fukaya | Future medicinal chemistry | 2015 | 10.4155/fmc.15.62 | https://pubmed.ncbi.nlm.nih.gov/26144262/ | cell signaling, oxidative stress, immunity, inflammation | organogermanium, Ge-132, cisdiol compounds, adrenaline, ATP, catecholamines, celltocell signaling, physiological functions, safety, antiinflammatory effects, antihypertensive effects, novel medical applications | Research objectives or hypotheses: Investigate interactions between Ge-132 and cis diol compounds like adrenaline and ATP, and their potential to regulate physiological functions<br>Setting and context: Conducted by researchers affiliated with the Asai Germanium Research Institute Co., Ltd., focusing on organogermanium compounds and their biological interactions | human cells keratinocytes | Number of cells per well: 6 × 10^5 cells/ml (100 μl per well)<br>Number of wells: Not explicitly mentioned<br>Statistical analysis: ANOVA with Tukey's multiple comparisons | Acute toxicity test: LD50 of 8500 mg/kg bodyweight in male dogs. Complex formation study: Equal moles of THGPA and adrenaline/noradrenaline (0.5 mol of Ge-132 in 10 ml water). Affinity studies: THGPA concentrations of 0.25 to 500 mmol/l. Cellular study: THGPA concentrations of 0, 2.5, and 5 mM for adrenaline; 0, 1, and 10 mM for ATP. | Ge-132 forms complexes with adrenaline and ATP; THGPA inhibits calcium influx caused by these compounds. Statistical results: Calcium ion influx significantly inhibited by 5 mM THGPA (p < 0.01); dose dependent inhibition by THGPA.Qualitative themes: Interaction between Ge-132 and cis diol compounds; potential for novel therapeutic applications. Significant relationships: High affinity of THGPA for catecholamines; inhibition of receptor binding by THGPA. Unexpected findings: Low toxicity of THGPA due to equilibrium state. Authors' interpretation: Ge-132 as a potential candidate for novel medicine through interaction with cis diol compounds. | THGPA modifies cis diol mediated cell to cell signaling; Ge-132 is a candidate for interacting with cis diols. Clinical implications: Potential novel targets for controlling cell cell communication. Practical implications: Ge-132 could lead to new medical applications. Recommendations: Further study on biological applications of organogermanium compounds. Limitations: High concentrations of THGPA needed for significant inhibition. Conflicts of interest: Authors employed by Asai Germanium Research Institute Co., Ltd. Generalizability: Potential applications across physiological systems. Overall assessment: Evidence suggests potential for Ge-132 in medical applications, but high concentrations required may limit effectiveness. | Japan |
| Eunhye Kim, Yubyeol Jeon, Dae Young Kim, Eunsong Lee, Sang-Hwan Hyun | Theriogenology | 2015 | 10.1016/j.theriogenology.2015.03.006 | https://doi.org/10.1016/j.theriogenology.2015.03.006 | Reproductive biology, oxidative stress, antiapoptosis | carboxyethylgermanium sesquioxide (Ge-132), antioxidative effect, in vitro maturation (IVM), porcine oocytes, parthenogenetic activation (PA), in vitro fertilization (IVF), glutathione (GSH), reactive oxygen species (ROS), embryonic development | Research objectives or hypotheses: Investigate the effect of Ge-132 on nuclear maturation, intracellular GSH and ROS levels, and subsequent embryonic development after PA and IVF<br>Setting and context: Use of porcine oocytes, application of Ge-132 during IVM, statistical analysis using SPSS 17.0 | porcine oocytes cells | Number of cells used in each experiment: Nuclear maturation: 967 oocytes. Intracellular GSH and ROS levels: 10 oocytes from each treatment group. Gene expression study: 160 mature oocytes and cumulus cells per group. Replicates: 3 per 3 times | 0 mg/mL (control), 100 mg/mL, 200 mg/mL, and 400 mg/mL Ge-132. | No significant difference in nuclear maturation among Ge-132 treatment groups. Significant increase in intracellular GSH levels and decrease in ROS levels with Ge-132 treatment. Higher cleavage rates and blastocyst formation rates in Ge-132 treated groups. Increased Nrf 2 mRNA expression and decreased Bax mRNA expression in Ge-132 treated oocytes. Improved developmental potential of PA and IVF porcine embryos due to enhanced antioxidant defenses and reduced oxidative stress. | Ge-132 improves developmental potential of porcine embryos by enhancing cytoplasmic maturation, increasing GSH levels, and reducing ROS levels and apoptosis. Clinical implications: Ge-132 could improve IVM and IVF efficiency. Recommendations: Further concentration experiments are needed. Study limitations: Potential toxic effects of high doses of Ge-132; need for further research on biological activities. Potential sources of bias or confounding: Not mentioned. Generalizability considerations: Not mentioned. Conflicts of interest or funding sources: Not mentioned. Overall assessment of evidence quality and reliability: Detailed experimental results and statistical analyses provided. | Republic of Korea |
| Yasuhiro Shimada, Katsuyuki Sato, Yoshihiko Tokuji, Takashi Nakamura | Carbohydrate research | 2015 | 10.1016/j.carres.2015.01.012 | https://doi.org/10.1016/j.carres.2015.01.012 | Biochemistry, Immunology, Diabetes Research | Nuclear magnetic resonance (NMR); Organic germanium compound Ge-132; Saccharides Monosaccharides; Glucose; Fructose; Cisdiol structures; Physiological functions; Complex formation | Research objectives or hypotheses: To evaluate the interactions between Ge-132 and monosaccharides using NMR spectroscopy<br>Setting and context: Using a Gemini 300 Spectrometer for NMR analysis, with samples dissolved in D2O and analyzed at specific concentrations and frequencies | in_vitro (water solution) | Not mentioned (the paper does not provide specific information on the number of replicate experiments or cells used) | 0.5 M for THGPNa when mixed with various monosaccharides (AHET, glucose, and fructose) | Ge-132 forms complexes with cis diol structures of saccharides. Variables measured: NMR spectra changes indicating complex formation. Qualitative themes: Higher affinity for fructose than glucose; b furanosyl conformation of fructose interacts most strongly. Significant relationships: Ge-132 interacts with specific saccharide structures; potential influence on glucose to fructose isomerization. Unexpected findings: Stronger interaction with fructose than glucose. Data tables: Table 1 shows ratio changes in NMR signals upon addition of D fructose. Authors' interpretation: Ge-132 may influence physiological functions and have industrial applications. | Ge-132 forms complexes with cis diol structures of saccharides, potentially implicating its physiological functions. Clinical implications: Ge-132 may interact with saccharides in vivo, affecting physiological functions.Theoretical implications: Ge-132 could influence isomerization from glucose to fructose. Recommendations: Further study on complex formation and interaction with various saccharides. Limitations: Need for further study to evaluate complex formation ratios. Potential sources of bias or confounding: Not mentioned. Generalizability: Findings may extend to oligo and polysaccharides. Conflicts of interest or funding sources: Not mentioned. Overall assessment of evidence quality and reliability: Detailed NMR spectroscopy data supports conclusions. | Japan |
| Takafumi Tezuka, Atsunori Higashino, Mitsuo Akiba, Takashi Nakamura | Advances in Enzyme Research | 2017 | 10.4155/fmc.15.62 | https://pubmed.ncbi.nlm.nih.gov/26144262/ | Antioxidant, Oncology, Inflammation | Organogermanium (Ge-132); Stress enzymes; Active oxygen species; Superoxide dismutase (SOD); Catalase (CAT); NADHdependent oxidase; NADPHdependent oxidase; Xanthine oxidase (XOD); Antioxidative effects; Physiological functions | Research objectives or hypotheses: To examine the effects of Ge-132 on the activities of superoxide forming and superoxide scavenging enzymes in a monkey liver preparation<br>Setting and context: In vitro study using a liver preparation from a Japanese macaque; context is the investigation of Ge-132's role in reducing active oxygen species levels | cells from liver of a Japanese macaque | Number of replicates: 3<br>Number of cells used per assay and condition: Not mentioned | NAD(P)H OD assay: 0, 0.001, 0.01, 0.1, or 1 μM Ge-132; SOD assay: 0, 0.001, 0.01, 0.1, or 1 μM Ge-132 CAT assay: 0, 0.001, 0.01, 0.1, or 1 μM Ge-132 | Ge-132 suppresses superoxide forming enzymes (NADH dependent oxidase, NADPH dependent oxidase, xanthine oxidase) and promotes superoxide scavenging enzymes (superoxide dismutase, catalase). Statistical results: Suppression of XOD by up to 90.8%, 89.1%, 80.8%, or 45.3% at different concentrations; promotion of SOD by up to 174.2%, 171.4%, 176.9%, or 177.5%; promotion of CAT by up to 162.9%, 182.2%, 188.7%, or 193.5%. Qualitative themes: Reduction in active oxygen species levels; potential protection against diseases like cancer, pulmonary complaints, rheumatism. Significant relationships: Ge-132 's role in maintaining low active oxygen species levels; potential health benefits. Data tables/figures: Figures 2 and 3 illustrate the suppression and promotion effects. Authors' interpretation: Ge-132 contributes to health improvement and disease protection by regulating stress enzymes. | Ge-132 suppresses stress enzymes and promotes superoxide scavenging enzymes, potentially reducing active oxygen species and improving disease recovery. Clinical implications: Potential use in protecting against diseases like cancer and rheumatism.Recommendations: Further application and research in medical and veterinary fields. Study limitations: Not explicitly mentioned. Potential sources of bias: Not mentioned. Generalizability: Implied applicability to various fields.<br>Conflicts of interest: None reported. Evidence quality and reliability: Supported by experimental data and lack of conflicts of interest. | Japan |
| Eunhye Kim, Seon-Ung Hwang, Junchul David Yoon, Eui-Bae Jeung, Eunsong Lee, Dae Young Kim, Sang-Hwan Hyun | The Journal of reproduction and development | 2017 | 10.1262/jrd.2017-020 | https://pubmed.ncbi.nlm.nih.gov/28993559/ | Antioxidant mechanisms, apoptosis, cell signaling pathways (KEAP1) | Carboxyethylgermanium sesquioxide (Ge-132); Oxidative stress; Apoptosis; In vitro culture (IVC); In vitro fertilization (IVF); KEAP1 pathway; porcine Embryonic development | Research objectives or hypotheses: To examine the effects of Ge-132 on porcine IVF embryos during IVC, focusing on reducing oxidative stress.<br>Setting and context: In vitro culture system for porcine embryos. | Porcine IVF embryos (zygotes and blastocysts) | Number of replicates: The experiment was repeated five times for some experiments, and at least three times for others.<br>Number of cells used per each assay and each condition: Detailed information is provided in tables, such as Table 1 for oocytes and Table 3 for embryos. | Ge-132 at 0, 100, 200, and 400 μg/ml. | Ge-132 treatment increased total cell numbers in blastocysts compared to controls. The 200 μg/ml Ge-132 group had significantly increased intracellular GSH levels and decreased ROS generation. mRNA expression levels of KEAP1 and proapoptotic genes (BAX and CASPASE3) were lower in Ge-132 treated blastocysts. Apoptotic and necrotic cell percentages were lower in Ge-132 treated embryos. KEAP1 protein was more localized in the cytoplasm of treated blastocysts. Blastocyst formation rates and hatched blastocyst percentages were higher in Ge-132 treated groups. Ge-132 activates an anti apoptotic pathway involving KEAP1 and BCL2, reducing oxidative stress and apoptosis. | Ge-132 improves embryonic development by reducing oxidative stress and apoptosis through the KEAP1 pathway. Clinical implications: Ge-132 could improve IVF success by reducing oxidative stress and apoptosis. Recommendations: Further investigation into Ge-132's mechanisms, especially its interaction with ATP. Limitations: Need for further experiments to understand Ge-132's mechanisms. Potential sources of bias or confounding: Not mentioned.Generalizability: Limited to porcine embryos; further research needed for other species. Conflicts of interest or funding sources: Not mentioned. Overall assessment of evidence quality and reliability: High due to detailed experimental design and statistical analysis. | Republic of Korea |
| Yasuhiro Shimada, Katsuyuki Sato, Tomoya Takeda, Yoshihiko Tokuji | BIOLOGICAL TRACE ELEMENT RESEARCH | 2018 | 10.1007/s12011-017-1020-4 | https://doi.org/10.1007/s12011-017-1020-4 | Biochemistry and molecular interactions, with a focus on anti-inflammatory activity and pain relief | Ge-132; THGP; Nucleic acid components; Adenosine deaminase (ADA); Antiinflammatory activity; Pain relief | Research objectives or hypotheses: Evaluate the interaction of THGP with nucleosides and nucleotides and assess its effect on adenosine deaminase activity.<br>Setting and context: Laboratory experiments using NMR spectroscopy and enzymatic assays. | in_vitro (water solution) | Number of replicates: At least three replicates for each condition.<br>Number of cells used: Not mentioned. | THGP: 0 100 mM Concentration of Ado or dAdo: 7.5 50 μM Concentration of ADA: 33.3 ng/mL in a final reaction volume of 600 μL | THGP forms complexes with nucleotides and nucleosides through their cis diol groups. The number of phosphate groups on the ribose moiety influences THGP's ability to form complexes. THGP inhibits ADA's catalysis of adenosine in a concentration dependent manner. Highest CFR value observed for Ado, followed by AMP, ADP, and ATP. K D values: Ado (28 mM), ATP (91 mM), nucleosides (23 ± 5 mM). THGP specifically inhibits ADA reactions involving adenosine due to its cis diol structure. Inhibitory effect on ADA reactions may minimize interference with cellular metabolism. Data tables provide detailed structures and K D values for nucleotides and nucleosides. | THGP interacts with nucleic acid components and inhibits ADA activity in a concentration dependent manner. Clinical implications: Potential use of Ge-132 to enhance pain relief by inhibiting ADA. Recommendations: Further studies to explore applications of Ge-132, particularly in reducing morphine or ADA inhibitor use. Study limitations: Not explicitly mentioned. Potential sources of bias: Not mentioned. Generalizability: Implied potential for medical applications.Conflicts of interest: None declared. Evidence quality: Strong experimental data and NMR analysis. | Japan |
| Takeyoshi Wada, Takashi Hanyu, Kota Nozaki, Kosuke Kataoka, Tomoro Kawatani, Toru Asahi, Naoya Sawamura | Biological and Pharmaceutical Bulletin | 2018 | 10.1248/bpb.b17-00949 | https://doi.org/10.1248/bpb.b17-00949 | Antioxidant activity, cell proliferation, immunity, oncology | Antioxidant activity, Ge-132, synthetic organic germanium, dietary supplement, cell proliferation, oxidative stress, hydrogen peroxide, immunological effects, antimetastatic mechanism | Research objectives or hypotheses: Investigate the cytotoxicity, cell proliferative effects, and antioxidant activity of Ge-132 on cultured mammalian cells<br>Setting and context: Use of CHO K1, HeLa, and SH SY5Y cells in DMEM medium with FBS at 37°C in 5% CO2 to examine effects on cell proliferation and oxidative stress induced by hydrogen peroxide | Cultured mammalian ovary cells; Chinese hamster ovary (CHO-K1) cells, HeLa cells, SH-SY5Y cells | Number of replicates: 3 to 5 independent experiments<br>Cytotoxicity assay: 1×10^5 cells per well in 12 well plates<br>Cell proliferation assay: 5×10^3 cells per well in 96 well dishes<br>Oxidative stress assay: 10×10^4 cells per well | Cytotoxicity assessment: 0 5 mM Ge-132 for 24 hours Cell proliferation studies: 0 5 mM Ge-132 Oxidative stress experiments: Pre incubation with 5 mM Ge-132 | Cytotoxicity (LDH levels), cell proliferation (ATP content), oxidative stress (ROS and carbonylated protein levels). No significant cytotoxic effect; significant increase in cell proliferation in CHO K1 and SH SY5Y cells; suppression of ROS production and reduction in carbonylated protein levels. Dose dependent effects on cell proliferation; antioxidant activity against hydrogen peroxide induced oxidative stress. Significant relationships or associations found: Ge-132 promotes cell proliferation and reduces oxidative stress. Unexpected or secondary findings: Potential neuroprotective effects suggested due to antioxidant activity. Data tables, figures, or charts: Figures 2, 3, 4, and 5 summarize key results. Authors' interpretation: Ge-132 has antioxidant activity and promotes cell proliferation, suggesting potential as an antioxidant supplement. | Ge-132 has antioxidant activity against hydrogen peroxide induced oxidative stress and promotes cell proliferation. Clinical implications: Potential neuroprotective effects. Recommendations: Future studies to identify neuroprotective effects and develop more effective Ge-132 derivatives. Study limitations: Unclear mechanism of antioxidant effect; need for further research on neuroprotective potential. Potential sources of bias or confounding: Not mentioned. Generalizability considerations: Limited to cultured mammalian cells; further in vivo studies needed. Conflicts of interest or funding sources: No conflict of interest declared. Overall assessment of evidence quality and reliability: Preliminary evidence from in vitro studies; requires further validation. | Japan |
| Galina E Filonova, Elena N Nikolaevskaya, Artem V Kansuzyan, Irina V Krylova, Mikhail P Egorov, Viatcheslav V Jouikov, Mikhail A Syroeshkin, N D Zelinsky | European Journal of Organic Chemistry | 2019 | 10.1002/ejoc.201900331 | https://doi.org/10.1002/ejoc.201900331 | Antioxidant properties | Antioxidant properties ;Adrenaline; Ge-132;Cyclic voltammetry; UV spectroscopy; 1 H NMR spectroscopy; Dietary supplement; Biochemical interactions | Research objectives or hypotheses: Investigate the effect of Ge-132 on the antioxidant activity of adrenaline in aqueous solutions<br>Setting and context: Study of complexation of Ge-132 with adrenaline in a phosphate buffer solution close to physiological conditions | rats tumor cells, ovary cells | Not mentioned (the paper does not provide information on the number of replicates or cells used per each assay and condition) | 1H NMR spectroscopy: Ge-132 at 1.3 × 10 1 M and diluted to 1.3 × 10 2 M Cyclic voltammetry: Ge-132 at 3 × 10 3 M, 6 × 10 3 M, and 9 × 10 3 M UV spectrophotometry: DPPH at 1 × 10 4 M; Ge-132 concentration not specified | The complex formed by adrenaline and Ge-132 is oxidizable at more positive potentials, indicating a more difficult oxidation process. The equilibrium constant for complex formation is estimated at 1300 M 1. At concentrations below 10 3 M, the complex is mostly dissociated, and Ge-132 's influence is significant only when in excess. The Ge-132 adrenaline complex has significantly reduced antioxidant properties compared to adrenaline alone. Dissociation at concentrations of 10 4 M or lower results in no detectable effect of Ge-132 on adrenaline's antioxidant properties. | : The complex of adrenaline and Ge-132 has reduced antioxidant properties compared to adrenaline alone. Clinical implications: Ge-132 may affect the biological activity of catechols like adrenaline. Recommendations: Careful consideration of Ge-132's impact on biological catechols is suggested. Study limitations: Findings are specific to studied concentrations and conditions. Potential sources of bias: Not mentioned. Generalizability: Limited to specific conditions and concentrations. Conflicts of interest: Not mentioned. Evidence quality: Supported by multiple spectroscopic methods. | Russia |
| Tomoya Takeda, Sota Doiyama, Junya Azumi, Yasuhiro Shimada, Yoshihiko Tokuji, Hiroaki Yamaguchi, Kosuke Nagata, Naoya Sakamoto, Hisashi Aso, Takashi Nakamura | Scientific Reports | 2019 | 10.1038/s41598-019-49883-7 | https://doi.org/10.1038/s41598-019-49883-7 | Antioxidant research, human dermal fibroblasts; cell signaling and inflammation; THGP | organogermanium, repagermanium, THGP, oxidative stress, normal human dermal fibroblasts, cell death, gene profiling, inflammation, skin protection, cosmetics | Research objectives: To evaluate the preventive effects of THGP against ROS induced NHDF death and to understand the mechanism through gene profiling and expression analysis.<br>Setting and context: Use of isotope microscopy to visualize germanium incorporation into NHDFs; gene profiling to understand THGP's protective effects against ROS induced cell death in the context of developing protective cosmetics. | normal human dermal fibroblasts (NHDFs) | MTS based measurement of cell proliferation: 8 replicates per condition, 1.5 × 10^4 cells/well.<br>LC MS/MS analysis: 4 replicates per condition, 2.0 × 10^5 cells/well.<br>Other assays: 1.5 × 10^5 cells/well for 12 well plates, 2.0 × 10^5 cells/well for six well plates. | In the cell viability assay under oxidative stress: 0, 5.9, 59, 590, and 5900 μM THGP. In the gene expression profiling: 0, 0.59, and 5.9 mM THGP. In the LC MS/MS analysis: 0, 1, and 10 mM THGP. In the PI/Hoechst staining: 0 and 5.9 mM THGP. | Reduction of cell death due to oxidative stress by THGP, suppression of genes related to cell death (NR4A2) and inflammation (IL6 and CXCL2), suppression of IL6 production. Statistical results: Significant downregulation of NR4A2, IL6, and CXCL2 genes; p < 0.05 for statistical significance. Qualitative themes or patterns identified: THGP's mechanism differs from general antioxidants; dose dependent effect on cell death suppression. Significant relationships or associations found: Suppression of cell death and inflammatory genes by THGP. Unexpected or secondary findings: THGP does not directly scavenge ROS. Data tables, figures, or charts: Tables on gene expression changes and statistical analysis results. Authors' interpretation: THGP suppresses cell death and inflammation through gene regulation, not ROS scavenging. | THGP act on gene and protein expression. Limitations: Unclear mechanism of THGP's effect on gene and protein expression. Potential sources of bias or confounding: Not mentioned. Generalizability: Potential applications in human skin care. Conflicts of interest or funding sources: No conflicts of interest declared; no funding sources mentioned. Overall assessment of evidence quality and reliability: Robust approach using multiple methods. | Japan |
| Yong-Sheng Wang, Guo-Qing Teng, Han Zhou, Chun-Liu Dong | BIOLOGICAL TRACE ELEMENT RESEARCH | 2020 | 10.1007/s12011-020-02106-x | https://doi.org/10.1007/s12011-020-02106-x | Inflammation, Immunity, Cell Signaling (NF-κB and MAPK pathways) | Germanium; Mastitis; Inflammation; NFκB pathway; MAPK pathway; TNFα; IL1β; IL6; IL10; Antiinflammatory effects; Clinical treatment | The study used a mastitis mouse model with sixty female BALB/c mice, divided into six groups. The study was conducted with ethical approval from the Institutional Animal Care and Use Committee of Jilin University. | In vivo: Mouse model (BALB/c mice)<br>In vitro: Primary mouse mammary epithelial cells (mMECs) | Total number of mice: 60<br>Number of mice per group: 10<br>Number of mMECs per condition: Not specified | In vivo experiments: Ge-132 at 5 mg/kg, 10 mg/kg, and 20 mg/kg, administered abdominally every 12 hours. In vitro experiments with mMECs: Ge-132 at 0.5 μg/mL, 1 μg/mL, and 2 μg/mL. | Histopathological changes, MPO activity, expression levels of TNF α, IL 1β, IL 6, IL 10. Statistical results: Ge significantly inhibited MPO activity and reduced TNF α, IL 1β, IL 6 expression; increased IL 10 expression in a dose dependent manner. Qualitative themes or patterns identified: Ge has a protective effect on mammary gland tissues by reducing inflammation. Significant relationships or associations found: Ge inhibits NF κB and MAPK pathway activation, reducing pro inflammatory cytokine expression. Authors' interpretation: Ge is an effective agent for treating mastitis by inhibiting inflammation. | Ge inhibits inflammation by inhibiting NF κB and MAPK pathways, potentially making it an effective treatment for mastitis and other inflammatory diseases. Clinical implications: Ge could be used to treat mastitis and other inflammatory diseases. Recommendations for future research: Development of drugs related to Ge could provide new treatments for inflammatory diseases. Study limitations: Not mentioned. Potential sources of bias or confounding: Not mentioned. Generalizability considerations: Not mentioned. Conflicts of interest or funding sources: Authors declare no conflict of interest. Overall assessment of evidence quality and reliability: Not explicitly mentioned. | China |
| Randall T Mertens, Sean Parkin, Samuel G Awuah | Inorganica chimica acta | 2020 | 10.1016/j.ica.2019.119375 | https://pubmed.ncbi.nlm.nih.gov/34565828/ | Oncology, Oxidative Stress | Germanium(IV)diketonate complexes; Anticancer agents; ROS accumulation; Mitochondrial membrane depolarization; Apoptosis induction; Synthesis using GeCl 2 •dioxane and acetylacetone; Anticancer activity in human cancer cell lines | Research objectives or hypotheses: Exploring the anticancer activity of germanium(IV) diketonate complexes and understanding their mode of action. Setting and context: Laboratory experiments using NMR, MS, X ray crystallography, cyclic voltammetry, and cell culture experiments | human cell lines MDA-MD-175, MDA-MB-231, MRC5, H460, A2780 | MDA MB 231 | Ge-132: 5 μM or 10 μM Duration: 48 hours for apoptosis analysis, 72 hours for cell cycle analysis, 4 hours for mitochondrial membrane potential assay. Stock solution: 5 mM or 10 mM in DMSO | These compounds show selective anticancer activity, with about threefold greater toxicity in cancer cells than in normal epithelial cells. They appear to act through a mechanism involving G1 cell-cycle arrest, ROS accumulation, and mitochondrial membrane depolarization, ultimately promoting apoptosis. In normal MRC5 cells, their IC50 values are around three times higher than those reported for cisplatin, suggesting lower toxicity toward non-tumoral cells. They are also stable in biological media, remain soluble in DMSO, and do not seem to be deactivated by biological nucleophiles such as GSH, NAC, or DNA bases. | Germanium(IV)-diketonate complexes show promise as anticancer agents with selectivity towards cancer cells and ability to induce apoptosis and ROS accumulation. Clinical implications: Potential use as anticancer agents with lower toxicity. Recommendations: Further development for cancer treatment, particularly for triple-negative breast cancer. Study limitations: Need for further ligand optimization; no significant difference in anionic effects.<br>Generalizability: Consistent results across multiple cancer cell lines. Evidence quality and reliability: High due to rigorous scientific methods. | USA |
| Sunanda Baidya, Yoko Nishimoto, Seiichi Sato, Yasuhiro Shimada, Nozomi Sakurai, Hirotaka Nonaka, Koki Noguchi, Mizuki Ki, Satoshi Tadano, Kozo Ishikawa, Kai Li, Aoi Okubo, Taisho Yamada, Yasu Orba, Michihi Sasaki, Hirofumi Sawa, Hiroko Miyamoto, Ayato Takada, Takashi Nakamura, Akinori Takaoka, Sakurai | Viruses | 2021 | 10.1128/MMBR.05015-11 | https://pubmed.ncbi.nlm.nih.gov/34578256/ | immunity, antiviral strategies, inflammation | Type I interferon (IFN) production; RIGI ligand 5'triphosphate RNA (3pRNA); Influenza virus (IAV); Vesicular stomatitis virus (VSV); Antiviral action; innate inflammation; Organogermanium compound THGP; RIGImediated viral sensing | Research objectives or hypotheses: To investigate the effects of THGP on RIG I mediated viral sensing and viral replication during influenza A virus infection.<br>Setting and context: Laboratory experiments using cell lines (RAW 264.7, HEK293T, A549, MDCK) and animal models (MAVS KO mice). | cell lines RAW 264.7, HEK293T, A549, MDCK) and animal (rats) | Cell experiments: 20,000 cells for fluorescence measurement; more than 30 cells for microscopic analysis<br>Animal experiments: Several mice (exact number not specified) | In vitro experiments: THGP concentration not explicitly mentioned, used in a dose dependent manner.<br>Animal experiments: Mice were pretreated with 30 μL of THGP (50 or 100 mg/kg) by i.n. injection at 3 h before infection. | THGP downregulates type I interferon production triggered by RIG-I ligands, especially 3pRNA.It competitively inhibits RIG-I recognition by binding the 5′-triphosphate group of viral RNA.THGP also directly inhibits influenza A virus replication by interfering with the interaction between viral polymerase and the viral RNA genome.This reveals a dual effect: modulation of antiviral immune signaling together with direct antiviral activity.The authors propose THGP as a potential therapeutic agent for RNA viral infections and inflammatory diseases. | THGP has a dual effect on suppressing RIG I mediated signaling and direct antiviral action against IAV, suggesting potential therapeutic implications. Clinical implications: THGP could be a therapeutic agent for IAV and possibly other viruses, as well as inflammatory diseases. Recommendations: Future research should explore the range of viruses affected by THGP. Limitations: THGP does not interact with viral RNAs of EMCV and SARS CoV 2. Potential sources of bias or confounding: Not mentioned. Generalizability: Limited by focus on specific viruses and mechanisms. Conflicts of interest: None declared. Overall assessment: The study provides novel insights into THGP's mechanisms, but further research is needed to fully explore its potential. | Japan |
| Junya Azumi, Yasuhiro Shimada, Tomoya Takeda, Hisashi Aso, Takashi Nakamura | International journal of molecular sciences | 2022 | 10.2131/fts.4.137 | https://pubmed.ncbi.nlm.nih.gov/36362152/ | inflammation | Inflammasome, Inflammation, ATP, THGP, Ge-132, IL1β, Caspase1, Pyroptosis, Metabolic diseases, Cardiovascular diseases, Inflammatory conditions, Neurological disorders | Research objectives: To investigate whether THGP suppresses inflammasome activation by forming a complex with ATP<br>Setting: Human THP 1 monocytes and RAW 264.7 murine macrophages were used | human cell THP-1 monocytes; mouse cells RAW264 | THP 1 cells: 24 well plate: 1 × 10^5 cells/well; 6 well plate: 1 × 10^6 cells/well; 96 well plate: 1 × 10^4 cells/well; RAW 264.7 cells: 24 well plate: 1 × 10^5 cells/well, Number of replicates not explicitly mentioned | THGP concentrations: 0.05 mM, 0.5 mM, 1.2 mM, 5 mM, 12 mM; Ge-132 concentration: 500 mM; ATP concentrations: 1 mM, 5 mM; BzATP concentration: 100 μM; NMR analysis concentration: 50 mM | IL 1β secretion, caspase 1 activation, pyroptosis, cell viability. Statistical results: THGP at 5 mM significantly inhibits IL 1β secretion by approximately 30%.<br>Qualitative themes or patterns identified: ATP dependent suppression of IL 1β secretion, suppression of pyroptosis.<br>Significant relationships or associations found: THGP's inhibitory effect on IL 1β secretion is stronger with ATP than with BzATP.<br>Unexpected or secondary findings: THGP may also suppress inflammasome activity in an ATP independent manner.<br>Authors' interpretation: THGP is expected to be a therapeutic/prophylactic drug for inflammasome related diseases. | THGP suppresses inflammasome activation by forming a complex with ATP, suggesting its potential as a therapeutic or prophylactic agent for inflammasome related diseases. Clinical implications: THGP could be used to treat or prevent diseases associated with excessive inflammasome activation. Recommendations: Further research is needed to examine the effects of THGP on inflammasome related diseases. Study limitations: The study did not examine the effects of THGP on inflammasome related diseases. Potential sources of bias or confounding: Not mentioned. Generalizability considerations: Not explicitly discussed. Conflicts of interest or funding sources: Not mentioned. Overall assessment of evidence quality and reliability: Not explicitly discussed. | Japan |
| F. Sekiguchi, Nene Koike, Yasuhiro Shimada, Kaho Sugimoto, Hiroshi Masuda, Takashi Nakamura, Hiroaki Yamaguchi, G. Tanabe, Shinsuke Marumoto, Yoshihito Kasanami, Maho Tsubota, T. Ohkubo, S. Yoshida, A. Kawabata | Redox biology | 2023 | 10.1016/j.redox.2022.102579 | https://pubmed.ncbi.nlm.nih.gov/36563535/ | Pain management, redox biology, calcium channel signaling and modulation | Ge-132; THGP; Sulfides; Cav3.2; Pain management; Inflammation; Cancer | Research objectives or hypotheses: To determine whether THGP can trap sulfide and inhibit sulfide induced enhancement of Cav3.2 activity, leading to suppression of Cav3.2 dependent pain<br>Setting and context: In vitro experiments using Cav3.2 transfected HEK293 cells and in vivo experiments using mice Data collection methods: Mass spectrometry and 1H NMR techniques; experiments on mice to assess pain responses | In vitro: Cav3.2-transfected HEK_293 (human kidney cell)<br>In vivo: Mice male_C57BL/6J, models of cyclophosphamide-induced cystitis and cerulein-induced pancreatitis, adult (≈8–10 weeks) | 1. In Vitro (HEK293 Cav3.2 patch clamp): Replicates: 4–5 different replicates per condition. 2. Exogenous sulfide pain model (in mice):n = 5 mice per group.3. Systemic administration (i.p.) pain reduction:Replicates: n = 10–12 mice per condition.4. Visceral pain models: in CPA induced cystitis:Replicates: n = 5–7 mice per condition in bladder assessments. In Cerulein induced pancreatitis:Replicates: n = 9 mice for pancreatic outcomes (weight, amylase). | 1. In Vitro (HEK293 Cav3.2 patch clamp):THGP concentrations tested: 1 mM, 3 mM, 10 mM.NaS challenge concentration: 10 μM.Measurements taken repeatedly every 12 s after stabilization for 10 min.Replicates: 4–5 different cells per condition (mean ± SEM).2. Exogenous sulfide pain model (in mice):Intraplantar $Na_2S$: 10 pmol/paw Local co injection (i.pl.) of THGP:0.02 μg/paw (≈ 0.1 nmol/paw)0.2 μg/paw (≈ 1 nmol/paw) 2 μg/paw (≈ 10 nmol/paw) Replicates: n = 5 mice per group (for A panel).3. Systemic administration (i.p.) pain reduction:THGP administered i.p. at:30 mg/kg ( ≈ 150 μmol/kg) 100 mg/kg (≈ 500 μmol/kg) Replicates: n = 10–12 mice per condition.4. Visceral pain models:CPA induced cystitis:Cyclophosphamide: 400 mg/kg i.p.THGP: 30 mg/kg and 100 mg/kg i.p. Comparator T channel inhibitor TTA A2 at 1 mg/kg. Replicates: n = 5–7 mice per condition in bladder assessments. Cerulein induced pancreatitis: Cerulein: 50 μg/kg i.p. ×6 injections (1 h interval) THGP: 100 mg/kg i.p. given 5 min after the final cerulein Replicates: n = 9 mice for pancreatic outcomes | Reduction in inflammation, pain, and cancer; inhibition of sulfide induced enhancement of Cav3.2 activity. Qualitative themes or patterns identified: THGP traps sulfide and inhibits its enhancement of Cav3.2 activity, leading to suppression of Cav3.2 dependent pain. Significant relationships or associations found: THGP reacts with sulfides to form a sulfur containing product; THGP inhibits sulfide induced enhancement of T type Ca2+ channel dependent membrane currents. Authors' interpretation: THGP traps sulfide and inhibits sulfide induced enhancement of Cav3.2 activity, leading to suppression of Cav3.2 dependent pain caused by sulfide applied exogenously and generated endogenously. | THGP traps sulfide and inhibits sulfide induced enhancement of Cav3.2 activity, leading to suppression of Cav3.2 dependent pain. Clinical, practical, or theoretical implications stated: THGP could be used to suppress pain related to sulfide induced enhancement of Cav3.2 activity. Recommendations for practice or future research: Not mentioned. Study limitations acknowledged by authors: Not mentioned. Potential sources of bias or confounding: Not mentioned. Generalizability considerations: Not mentioned. Conflicts of interest or funding sources: Not mentioned. Overall assessment of evidence quality and reliability: Not mentioned. | Japan |
| Junya Azumi, Tomoya Takeda, Yasuhi Shimada, Tao Zhuang, Yoshihiko Tokuji, Naoya Sakamoto, Hisashi Aso, Takashi Nakamura | International journal of molecular sciences | 2023 | 10.3390/ijms24031885 | https://pubmed.ncbi.nlm.nih.gov/36768216/ | Immunity, Oncology | Organogermanium; THGP; M1 Macrophages; Tumor Immunology; Phagocytosis; Melanoma; NFκB Activation; SIRPα; CD47; Cancer Therapy; Immunomodulation | Research objectives: Investigate the effect and mechanism of THGP in inducing M1 macrophage differentiation and suppressing melanoma cell proliferation in vitro and in vivo.<br>Setting and context: In vitro experiments using RAW 264.7 cells and in vivo studies using mice, conducted at Asai Germanium Research Institute Co., Ltd., Japan. | RAW 264.7 mouse macrophage-like cells | Not mentioned (the paper does not provide the number of replicates for each assay or condition) | In vitro experiments: RAW 264.7 cells were cultured with 500 μM THGP. Dose dependent effect: THGP concentrations ranged from 50 to 5000 μM. In vivo experiments: Mice were fed diets containing 0.05% or 0.01% Ge-132. NF κB inhibitor: JSH 23 was used at 5 μM. | Macrophage differentiation into M1 phenotype, suppression of melanoma cell proliferation, expression of M1 and M2 markers, phagocytic activity.Statistical results: THGP increased M1 markers (iNOS by 1.38 fold, CD80 by 2.23 fold, CD86 by 2.23 fold, TNF α by 1.32 fold, IL 1β by 2.81 fold) and decreased M2 markers (arginase by 0.83 fold, CD206 by 0.34 fold, CD163 by 0.69 fold, TGF β by 0.64 fold, CXCL2 by 0.15 fold). Qualitative themes or patterns identified: THGP promotes M1 polarization and suppresses SIRP α and CD47 expression, enhancing phagocytosis.Significant relationships or associations found: NF κB activation is crucial for M1 differentiation; suppression of SIRP α and CD47 enhances phagocytosis. Unexpected or secondary findings: THGP does not affect the production of cytotoxic cytokines but increases phagocytic ability. Data tables, figures, or charts: Figures 1 7 provide visual summaries of key results. Authors' interpretation: THGP is a potential anticancer agent by promoting M1 macrophage differentiation and enhancing phagocytosis. | THGP induces M1 macrophage differentiation, enhancing phagocytic activity and cytotoxicity against melanoma cells. Clinical implications: THGP could be a new regulatory agent for suppressing tumor immunity. Recommendations: Further research on THGP's mechanism and potential as an anticancer agent. Limitations: Unclear mechanism behind THGP's effect on SIRP α expression. Potential biases: Conflicts of interest due to funding and author affiliations. Generalizability: Limited by specific cell lines and animal models. Conflicts of interest: Authors affiliated with Asai Germanium Research Institute Co., Ltd., which funded the study. Evidence quality: High due to detailed experimental design, but potential biases should be considered. | Japan |
| Tairini E Cruz, \| Luan, Sitó-Silva, \| A Rogério, Almeida Filho, \| Alicio, Martins Júnior, Fernanda N Marqui, Diego G Souza, Tatiana I U Berton, Camila P Freita-Dell'aqua, Eunice Oba | Reproduction in domestic animals = Zuchthygiene | 2024 | 10.1111/rda.14703 | https://pubmed.ncbi.nlm.nih.gov/39149931/ | Antioxidant effects on bovine semen cryopreservation; ROS; Ge-132 | Antioxidant; Biscarboxyethyl germanium sesquioxide (Ge-132); Bovine semen cryopreservation; Morphofunctional sperm parameters; Sperm motility; Sperm membrane integrity; Reactive oxygen species; Lipid peroxidation; Sperm viability | Research objectives: Investigate the effects of Ge132 on sperm motility and morphofunctional integrity, as well as reactive oxygen species production<br>Setting and context: Conducted at the Tairana Artificial Insemination Centre in Presidente Prudente, São Paulo, Brazil, with ethical approval from the National Council for Animal Experimentation Control and the Ethics Committee for the Use of Animals | bovine sperm cells | Number of cells used per each assay: 10,000 cells for flow cytometry *3 replicates. Number of cells used per each condition: Diluted to 5 × 10^6 spermatozoa/mL for flow cytometry | 0 μg/mL , 500 μg/mL ,1000 μg/mL of Ge132 incubated at 33°C for 30 min; 0 μg/mL, 500 μg/mL ,1000 μg/m of Ge132 immediately cooled to 4°C | Total motility (TM), progressive motility (PM), curvilinear velocity (VCL), progressive velocity (VSL), average path velocity (VAP), amplitude of lateral head displacement (ALH), beat cross frequency (BCF), linearity (LIN), straightness (STR), lipid membrane peroxidation (LMP). Statistical results: TM higher in R0 compared to D0, D500, and R1000; significant differences in ALH, LIN, and STR among treatment groups. Qualitative themes: Ge132 may have beneficial effects on sperm kinetics; pre incubation with Ge132 maintains sperm motility and potentially inhibits early sperm capacitation. Significant relationships: Increased LIN and STR in Ge132 treated groups; reduced lipid peroxidation with Ge132. Unexpected findings: Ge132 did not significantly alter total motility; antioxidant effect not strongly evident. Data tables: Tables 1, 2, and 3 provide detailed results on sperm parameters and lipid peroxidation. Authors' interpretation: Ge132 may contribute to fertilization potential by improving kinetic parameters despite not strongly altering total motility. | The antioxidant effect of Ge132 was not clearly evident, but it may contribute to fertilization potential by improving sperm kinetics. Clinical implications: Ge132 may have some benefits for sperm kinetics, but its overall impact on fertility is unclear.Recommendations: Further research is needed to understand metabolic activity and to clarify the effects of Ge132. Limitations: The study had difficulty in understanding metabolic activity and its consequences. The presence of other antioxidants and temperature conditions may have affected results. Potential sources of bias: Presence of other antioxidants in the freezing medium and temperature conditions. Generalizability: Limited by specific conditions and concentrations used.Conflicts of interest: Donations from Botupharma Biotecnologia Animal and support from Central de Inseminação Artificial Tairana. Evidence quality: The study's findings are not strongly conclusive due to limitations and potential confounding factors. | Brazil |
| Tomoya Takeda, Junya Azumi, Mika Masaki, Takae Nagasawa, Yasuhiro Shimada, Hisashi Aso, Takashi Nakamura | Heliyon | 2024 | 10.1016/j.heliyon.2023.e23296 | https://doi.org/10.1016/j.heliyon.2023.e23296 | Immunology, Hematopoiesis, Antioxidant properties, Cardiovascular health, Exercise performance, Gastrointestinal health | Ge-132, RBC clearance, macrophage activation | Research objectives: Evaluate the effect of THGP on RBC phagocytosis by macrophages, investigate metabolic pigments in feces, and examine the effect of Ge-132 on erythroblast differentiation.<br>Setting and context: Male C57BL/6 J mice and male ICR mice, RAW264.7 mouse macrophage like cells.Laboratory experiments with controlled environmental conditions and statistical analyses using Student's t test and Dunnett's and Steel's methods. | animal (rats C57BL/6 and ICR) and in_vitro (cell lines RAW264.7) | In vitro experiments: 5 × 10^5 RAW264.7 cells per well, n = 5 6 replicates.<br>Animal experiments: First experiment: 2 groups of 6 mice each. Second experiment: 2 groups of 8 mice each. Third experiment: 3 groups of 10 mice each. | In vitro experiments with RAW264.7 cells: 0, 50, 500, and 5000 μM THGP.In vivo experiments with mice: 0.05% Ge-132 in diet for 4 or 7 days. | THGP increased phagocytic activity of RAW264.7 macrophage like cells. Ge-132 consumption changed fecal color to bright yellow, indicating increased erythrocyte degradation. Ge-132 increased erythrocyte metabolic pigments and antioxidant activity in feces. Hematocrit levels remained normal despite increased erythrocyte degradation. Ge-132 promoted erythroblast differentiation in bone marrow cells. Antioxidant capacity in feces was significantly increased with Ge-132 ingestion. GSH/GSSG ratio was increased in erythrocytes from Ge-132 fed mice. BFU E colonies were significantly increased in Ge-132 fed mice | Ge-132 promotes the clearance of senescent red blood cells via macrophage mediated phagocytosis, enhancing RBC degradation and production. Clinical implications: Potential improvements in blood flow and oxygen transport. Limitations: Study conducted in healthy mice and cell lines ; further research needed for human application. Conflict of interest: Financial relationships between authors and Asai Germanium Research Institute Co., Ltd. Generalizability: Needs further investigation for human health and disease conditions | Japan |

| | | | |
|---|---|---|---|
| Antioxidant | NRF2 | ATGACAATGAGGTTTCTTCGG | FW |
| | | CAATGAAGACTGGGCTCTC | REV |
| | HMOX1 | AACTCCCTGGAGATGACTC | FW |
| | | CTCAAAGAGCTGGATGTTGAG | REV |
| | GCLM | GTTGACATGGCCTGTTCAG | FW |
| | | AACTCCATCTTCAATAGGAGGT | REV |
| | GCLC | AAGTGGATGTGGACACCAG | FW |
| | | CTGTCATTAGTTCTCCAGATGC | REV |
| | NQO1 | ACATCACAGGTAAACTGAAGG | FW |
| | | TCAGATGGCCTTCTTTATAAGC | REV |
| MACROAUTOPHAGY | SQSTM1 | CTGGGACTGAGAAGGCTCAC | FW |
| | | GCAGCTGATGGTTTGGAAAT | REV |
| | NDPS2 (CALCOCO2) | ACCATGGAGGAGACCATCAA | FW |
| | | TTCTGGACGGAATTGGAAAG | REV |
| | ULK1 | TCATCTTCAGCCACGCTGT | FW |
| | | CACGGTGCTGGAACATCTC | REV |
| | ATG2B | AACTCACAAACAGAATGGTTCAAA | FW |
| | | AAGGGTACCAGGAAGACACCA | REV |
| | ATG3 | CATGCAGGCATGCTGAGGTG | FW |
| | | CGTTAACAGCCATTTTGCCACT | REV |
| | ATG4D | CACATCCTCAGGAAAGCCGT | FW |
| | | GACCACAGACTTCCACTCGG | REV |
| | ATG5 | GGGAAGCAGAACCATACTATTG | FW |
| | | AAATGTACTGTGATGTTCCAAGG | REV |
| | ATG7 | AGGAGATTCAACCAGAGACC | FW |
| | | GCACAAGCCCAAGAGAGG | REV |
| | GABARAPL1 | ACCATGGGCCAACTGTATGA | FW |
| | | TGGGCTTCCAACCACTCATTT | REV |
| LYSOSOMES | LAMP2A | GTGCAACAAAGAGCAGACTGT | FW |
| | | GGCACAAGGAAGTTGTCGTC | REV |
| | LAMP2B | AGAGTGTTCGCTGGATGATG | FW |
| | | TGCCAATTACGTAAGCAATCA | REV |
| | LAMP2C | AAGGGTTCAGCCTTTCAATG | FW |
| | | ACAATTATAAGGAAGCCCAAGG | REV |
| INFLAMMATION | p53 | CCTCAGCATCTTATCCGAGTGG | FW |
| | | TGGTGAGGATGGGCCTCC | REV |
| | IL-6 | AAATTCGGTACATCCTCGACG | FW |
| | | TTTCACCAGGCAAGTCTCC | REV |
| | TNFa | CTCTTCTGCCTGCTGCACTTTG | FW |
| | | CCACTGGAGCTGCCCCTCAG | REV |
| | Tert-1 | AGAGTGTCTGGAGCAAGTTG | FW |
| | | AGTCCATGTTCACAATCGGC | REV |
| | NFκb | GAACCACACCCCTGCATATAG | FW |
| | | GCACAACTCCTTCATCCTCTC | REV |
| PRE-GLYCATION | GLO1 | ATTGCCGTTCCTGATGTCTAC | FW |
| | | GAATCTCAATCCAGTAGCCGTC | REV |
| | GLO2 | TGGGCTGAAGGTTTATGGAG | FW |
| | | CAGATGTGTCCCGAAGTATGG | REV |
| | DJ-1 | ATTTATCTGAGTCGCCTATGGTG | FW |
| | | GCATCCAAAACCTACTTCGTG | REV |
| Housekeeping | BETA ACTIN | GTACGACCAGAGGCATACAG | REV |
| | | ACCGTGAAAAGATGACCCAG | FW |
